# UBE3C links ubiquitin signaling to epitranscriptomic control of cortical neurogenesis

**DOI:** 10.1101/2025.04.09.646620

**Authors:** Ekaterina Borisova, Katherine J. Cuthill, Rike Dannenberg, Janina Koch, Julius Nowaczyk, Theres Schaub, Manuela Schwark, Nicolai Kastelic, Ivanna Kupryianchyk-Schultz, Marieluise Kirchner, Tancredi Massimo Pentimalli, Thornton J. Fokkens, Frank Stein, Carola Dietrich, Claudia Quedenau, Tatiana Borodina, David Schwefel, Thomas Conrad, Agnieszka Rybak-Wolf, Nikolaus Rajewsky, Philipp Mertins, Per Haberkant, Sonja Lorenz, Nils Brose, Hiroshi Kawabe, Victor Tarabykin, Mateusz C. Ambrozkiewicz

**Author notes:** Senior authors (Nils Brose, Hiroshi Kawabe, Victor Tarabykin, Mateusz C. Ambrozkiewicz). These authors contributed equally (Ekaterina Borisova, Katherine J. Cuthill, Rike Dannenberg).

## Abstract

Neurological conditions are the leading cause of ill health worldwide. Here, we show that the neurodevelopmental disorder-associated ubiquitin ligase UBE3C regulates the cellular composition of the murine cerebral cortex and human brain organoids, with its loss favoring neurogenesis and suppressing glial fate. Using genetic complementation, we demonstrate that disease-associated *UBE3C* mutations alter its autoubiquitination activity and disrupt cortical lamination. Proteomic profiling of *UBE3C*-deficient forebrains and organoids identifies Cbll1 as a UBE3C substrate, and we show that the UBE3C-Cbll1 duo drives N^6^-methyladenosine (m6A) mRNA methylation. Hyperactivation of m6A writers in *UBE3C*-deficient neural progenitors impairs cell cycle exit, a defect reversible in vivo by the METTL3 inhibitor STM2457. Our findings uncover an epiproteomic mechanism controlling m6A-mediated gene expression and define a regulatory axis linking ubiquitin signaling to epitranscriptomic control of neural fate. This work provides a mechanistic framework for understanding neurodevelopmental disorders and highlights potential therapeutic strategies.

## INTRODUCTION

Approximately one in ten children is diagnosed with a neurodevelopmental disorder (NDD), which affects emotional, social, and cognitive development from an early age and persists throughout life ^1^. Developmental assessments are often sought for conditions such as global developmental delay, autism spectrum disorder (ASD), and specific learning disabilities. From an evolutionary perspective, the cerebral cortex represents the most recent evolutionary advance in our brain and serves as the biological foundation for our cognitive abilities ^2^. Modern human-specific cognitive adaptations are correlated with the enlargement of the neocortex ^3^. However, absolute brain size alone is insufficient to explain mammalian cognition, as the evolution of intelligence has also been shaped by brain morphology, the relative expansion of higher-order cortical areas, and the intricate connectivity between neuronal types ^4,5^. It is becoming increasingly clear that the origins of NDDs are associated with early molecular changes observed in neuronal stem cells ^6^. Such alterations affect the cellular composition of the adult brain, its connectivity, and the physiology of neuronal networks.

A vast majority of glutamatergic neurons in the mammalian cortex derive from ventricular progenitor cells, which retain the ability to self-renew and through their divisions, specify neurons and glia ^7,8^. Upon radial migration and termination in one of the six cortical layers, newly born neurons extend their axons and dendrites to establish synapses and form the framework of functional circuits ^9–11^. After neurogenesis ceases, cortical progenitors give rise to astrocytes ^12^. Disruptions to any of the developmental milestones during corticogenesis have been reported for NDD-associated genes ^13^. One of the prominent features of the cerebral cortex is the remarkable diversity of its cell types originating from neuronal stem cell niche, which specify stereotypic cellular lineages over developmental time ^14^. One of the core disease mechanisms disrupted in NDDs is aberrations in the composition of cellular lineage specified by cortical progenitors, resulting in either fewer neurons populating the cortex and primary microcephaly ^15^, or an imbalance in the proportion of cortical neuron subtypes altering the cortical efferents, as reported for ASD ^16^. Specification of cortical cell fate "cost" is embedded within cortical progenitors, whereby precocious acquisition of one cellular fate may diminish the subsequently specified cell type, representing an inherent molecular zero-sum game ^17–19^.

For decades, unraveling the molecular factors driving cellular diversification in the neocortex has remained a central focus in developmental neuroscience. In recent years, excellent research has delineated the transcriptomes of single cells in the mammalian cortex, generating comprehensive maps and atlases of cellular states and differentiation trajectories ^20–23^. In our recent work, we have shown that on top of elegant transcriptional programs and metabolic states ^24,25^, (post-)translational regulation of gene expression is indispensable for guiding fate decisions in neural stem cells, orchestrating cerebral cortex assembly in the developing brain ^17,26,27^. Regulating the stability, turnover, and protein functions in a cell-type-, cell-compartment-, and physiological-state-specific manner, orchestrated protein degradation leveraging the ubiquitin-proteasome system is among the principal signaling modes that make up the molecular logic of neurodevelopment ^27–29^. Ubiquitination comprises a multilayered biochemical machinery with E1 activating, E2 conjugating enzymes, E3 ubiquitin ligases, deubiquitinases, and a robust ubiquitin code ^30^, as well as an ever-growing molecular repertoire of ubiquitinated molecular species ^31^. The critical role of ubiquitin-mediated proteostasis is particularly evident in the developing brain, as approximately 13% of all E3 ligases are directly linked to neurological diseases, with NDDs at the forefront ^32,33^.

In this project, we investigated the role of Ubiquitin Protein Ligase E3C, UBE3C, which belongs to the Homologous to E6-AP C-terminus (HECT)-type family of E3 ligases. Loss-of-function mutations in *UBE3C* are associated with biallelic rare NDD with clinical diagnosis resembling the Angelman syndrome. Neurological diagnosis of *UBE3C* null patients include motor delay, severe intellectual disability (ID), limb dystonia, mutism, and neurobehavioral issues, such as aggression, overeating, and hyperactivity ^34^. *De novo* missense variants in *UBE3C*, leading to single amino acid substitutions, such as S845F ^35^ and F996C ^36^, have been found in patients with sporadic autism. Although UBE3C has been studied in other principal tissues ^37,38^, its role in the developing brain, especially regarding its protein substrates and the molecular etiology of NDD and ASD remains unclear.

Here, we find that NDD- and ASD-associated UBE3C orchestrates the output of cortical progenitors and the formation of cortical upper layers. Using biochemical assays with recombinant variants of UBE3C, we show that ASD-associated point mutations alter enzymatic activity and thermal stability of the enzyme and fail to restore cortical neuron lamination in genetic complementation experiments using conditional knock-out (cKO) *Ube3C* mouse line in vivo. Leveraging Tandem Ubiquitin Binding Entitites (TUBE) proteomics, single-nuclei RNA sequencing (snRNAseq) and bromodeoxyuridine- (BrdU-) birth-dating, we demonstrate that the Ube3C ubiquitome converges on the regulation of the mitotic cycle in cortical progenitors. Loss of *UBE3C* in murine cortical primordium and human cerebral organoids causes an overproduction of neurons at the expense of astrocytes by protracting neurogenic divisions of cortical progenitors and premature activation of differentiation programs for neuronal lineages. Differential expression analysis using quantitative proteomics in the neocortex identifies Cbl Proto-Oncogene Like 1, Cbll1, as a Ube3C substrate regulating cortical lamination and the composition of progenitor lineages. Next, using proteomics-based interactome identification, we show that Ube3C-Cbll1 duo associates with METTL Associated Complex (MACOM) to positively drive N^6^-methyladenosine (m6A) writing activity in the embryonic neocortex. Finally, we reveal that pharmacologically inhibiting the methyltransferase Mettl3 restores aberrant progenitor cell cycle exit in *Ube3C* KO mice. Taken together, our study defines the molecular etiology of UBE3C-associated NDDs, reveals a pharmacologically tractable pathway, and further expands the many roles of the ubiquitin signaling in the developing brain. By establishing UBE3C as a regulator of m6A RNA methylation through targeted degradation of Cbll1, we extend the ubiquitin code into the realm of epitranscriptomic control. These findings broaden the mechanistic landscape of neural progenitor domain beyond transcriptional and chromatin-based regulation, introducing a new axis of post-translational-post-transcriptional crosstalk.

## RESULTS

### Generation of cortex-expressed *UBE3C* knock-out models

UBE3 ligases, including UBE3A and UBE3B are linked to severe neurological diseases in human patients, reflecting neuron-specific molecular roles of these enzymes ^28,39^. Because of the association between mutations in *UBE3C* gene with patients suffering from severe NDD and ASD, we hypothesized that the UBE3C E3 ligase has fundamental roles in the developing brain.

First, we corroborated that its murine ortholog, Ube3C, was expressed throughout the entire corticogenesis, with higher levels towards the end of murine neurogenesis at embryonic day (E) 15.5 – 16.5, as shown by Western blotting (Fig. S1a). Using fluorescence in situ hybridization (FISH), with specific anti-Ube3C probe, we detected pan-cortical Ube3C expression throughout development in critical cellular niches (Fig. S2), including the ventricular zone (VZ) and the cortical plate (CP).

NDD patients with severe neurological symptoms possess biallelic loss-of-function mutations in *UBE3C* ^34^, whereas ASD patients carry point substitutions ^35,36^. To genetically knock-out (KO) *Ube3C* in mice, we took advantage of Cre-*loxP* technology. To circumvent the possibility of overt phenotypes plausibly present in *Ube3C* ^-/-^ ^28,34^, we decided to study the role of Ube3C in the developing neocortex using *Ube3C* ^f/f^ mice and conditional *Emx1*^Cre/+^ line (Fig. S1b-S1e). This driver allows for the deletion of *Ube3C* in dorsal telencephalic progenitors from E9.5 onward ^40^. Such an approach enables conditional KO (cKO) of *Ube3C* in the glutamatergic neurons and astrocytes of the cerebral cortex and hippocampus and allows us to dissect the cell-autonomous roles of Ube3C. Using an antibody against Ube3C, we corroborated the loss of Ube3C protein in the cKO brains of postnatal day (P) 0 mice (Fig. S1a), validating our targeting strategy. The prevailing expression of Ube3C in the cKO sample was associated with the presence of non-*Emx1*-lineage cells, such as interneurons or microglia.

Ube3C belongs to the class of HECT-type ubiquitin ligases. Prior to the transfer of ubiquitin to substrate proteins, the catalytic Cys residue in the C-terminal portion of the catalytic HECT domain forms a thioester-linked complex with ubiquitin (Fig. S1f) ^41,42^. To study Ube3C biochemically, we generated the HECT domain of Ube3C (HECT^Ube3C^) and its dominant negative variant C1051S, which is unable to accept ubiquitin. We validated the activity of both HECT variants in a reconstituted autoubiquitination assay, which monitors ubiquitin conjugation in the presence of ATP, Mg^2+^, E1 activating, and E2 conjugating enzymes (Fig. S1g). The exclusion of the E1 activating enzyme, or the substitution of the critical Cys residue to Ser abrogates the reaction, whereas wild-type HECT^Ube3C^ assembled robust polyubiquitin, as visualized by Coomassie staining (Fig. S1g).

Finally, we used the CRISPR-Cas9 system to delete *UBE3C* in human induced pluripotent stem cells (iPSCs). We targeted exon 7 of *UBE3C* using a single sgRNA, given its high on-target efficacy and low off-targeting in silico. Clones were verified by Sanger sequencing (Fig. S1h) and Western blotting (Fig. S1i) and were able to differentiate into forebrain organoids (Fig. S1j). We generated an isogenic control line and two *UBE3C* KO iPSC lines.

### Ube3C is indispensable for the formation of cortical upper layers

Given the severe NDD and ASD observed in patients with mutations in *UBE3C*, we first asked whether Ube3C influences the development of upper layer cortical neurons, harboring higher-order processing brain centers and responsible for interhemispheric consolidation, fundamental for cognitive abilities ^43^.

We used in utero electroporation (IUE), a method of in vivo DNA delivery into the dorsal ventricular progenitors of the cortex at E14.5, the onset of upper layer neurogenesis ^17,27^. As predicted from their birth date, four days later, at E18.5, the majority of control neurons localized correctly at the top of the CP (Fig. 1a). Expression of a catalytically inactive point mutant unable to accept ubiquitin (Fig. S1f-S1g), C1051S, resulted in a loss of upper layer localization of targeted neurons (Fig. 1b-1d; compare Fig. S3a for the uncropped IUE images), indicative of the requirement of Ube3C enzymatic activity for proper formation of upper layers during corticogenesis.

**Fig. 1.**
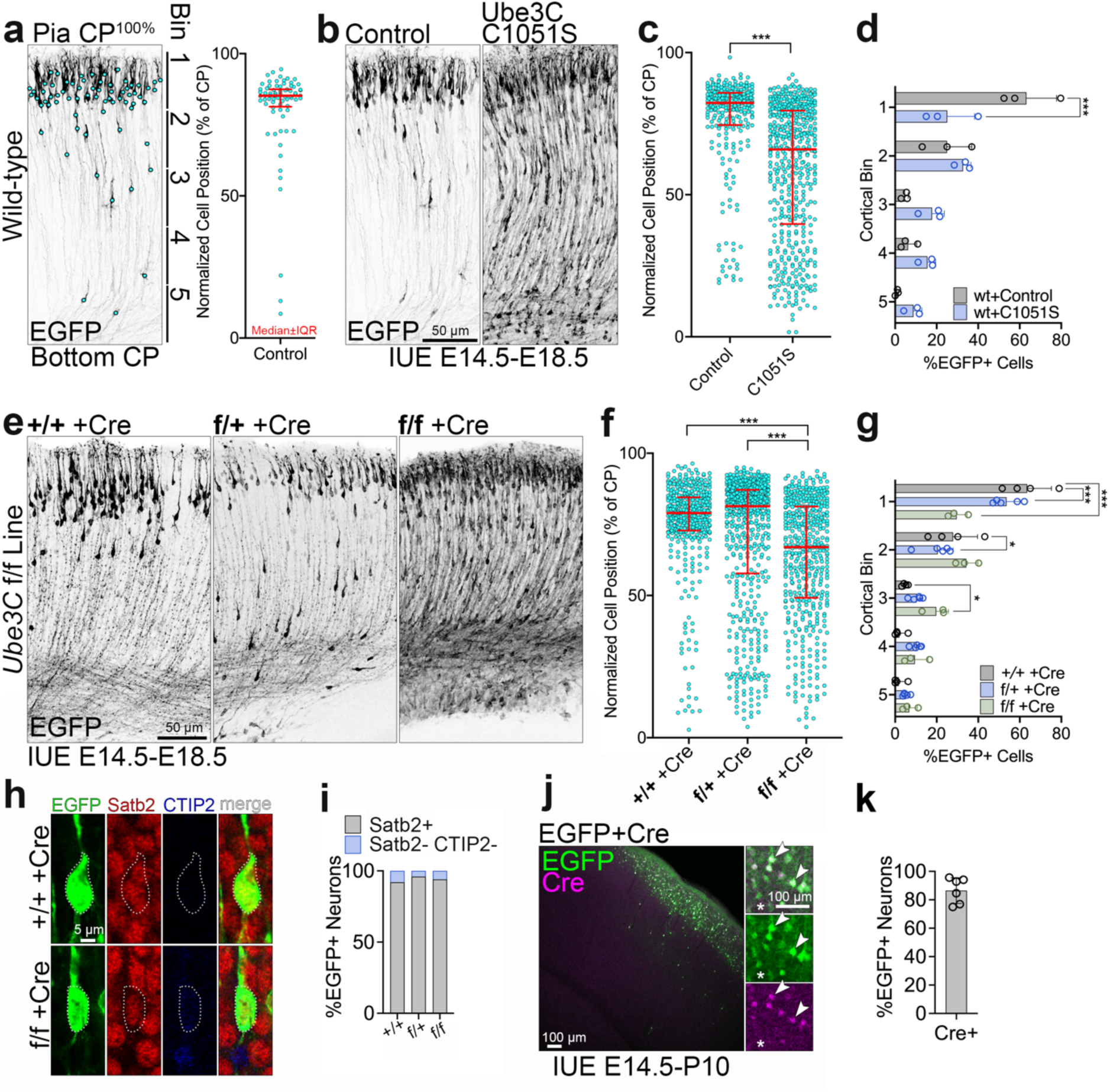
Ube3C is indispensable for proper laminar positioning of upper layer neurons. (a) The analysis of laminar positioning of neurons after IUE. Left: EGFP immunostaining signals in E18.5 50 μm-thick coronal cortical sections from wild-type embryos after IUE at E14.5 with a plasmid encoding for EGFP and manual markings of somata positions. Right: quantification of neuronal distribution across the CP. The position of each neuron was normalized to the thickness of the CP. 0% - bottom of the CP, 100% - pia. Graph contains pooled data from indicated number of brains. (b) EGFP immunostaining in coronal cortical sections from wild-type embryos after IUE at E14.5 with plasmids to express equimolar amounts of EGFP (control) or EGFP and Ube3C C1051S. (c-d) Neuronal distribution for the experiment in (b). (d) Laminar positioning of cortical neurons, in cortical bins, normalized to the global IUE efficiency. (e) EGFP immunostaining in coronal cortical sections from embryos of indicated *Ube3C* genotypes after IUE at E14.5 with plasmids to co-express EGFP and Cre. (f-g) Neuronal distribution after IUE described in (e). (h) Example results of immunolabeling for fate markers in (e). Somata are outlined with dotted lines. (i) Quantification of marker expression in electroporated cells after (e). No CTIP2-positive cells were found. (j) EGFP and Cre immunostaining in coronal cortical section after IUE at E14.5 to co-express EGFP and Cre. Arrowheads point to double positive neurons and the star marks a Cre-negative electroporated neuron. (k) Quantification of co-transfection efficiency by IUE using two plasmids described in (j). Bar graph and error bars on (d), (g), and (k) depict average and S.D. Line and error bars on (a), (c) and (f) indicate median and interquartile range (IQR). Number of brains analyzed for (c-d), and (f), n CTR =3, n C1051S =3; for (e-g), n +/+ =4, n f/+ =5, n f/f =3; for (i), n +/+ =74, n f/+ =90, n f/f =154 cells from two brains per genotype; for (k) n=6 fields of view from two brains. Statistics for (c), and (f), D’Agostino-Pearson normality test and Mann-Whitney test (c), or Kruskal-Wallis with Dunn’s multiple comparisons test (f); (i), Chi-square test; for (d), and (g), two-way ANOVA with Šidák multiple comparisons test. *** p < 0.001; * 0.01 < p < 0.05.

We then studied the consequence of *Ube3C* loss on upper layer neuron lamination. We used IUE to co-transfect EGFP and Cre-encoding plasmids into E14.5 progenitors of *Ube3C* ^+/+^, ^f/+^ and ^f/f^ embryos, thereby inducing *Ube3C* KO in single neuronal lineages (Fig. 1e). Consequently, the loss of *Ube3C* leads to an aberrant laminar distribution of upper layer neurons, as compared to properly localizing control ones. Interestingly, loss of a single *Ube3C* allele resulted in a moderate lamination defect, indicative of *Ube3C* dose dependence (Fig. 1e-1g and Fig. S3b). Lamination defects were independent of the neuronal fate, with the majority of E14.5 progenitor-born neurons expressing Special AT-Rich Sequence-Binding Protein 2, Satb2 ^10,17^ (Fig. 1h-1i and Fig. S3b).

Co-transfection of plasmids encoding EGFP and Cre into E14.5 progenitors led to co-expression of both proteins in the vast majority of neurons at P10 (Fig. 1j-1k). These results indicate that Ube3C activity is critical for the correct positioning of properly specified upper layer neurons within the developing CP.

### ASD-associated point mutations alter the thermodynamic properties of Ube3C and fail to restore proper cortical neuron lamination

Having established that genetic inactivation of *Ube3C*, mimicking the scenario in NDD patients, hinders the organization of the CP into layers, we next evaluated the properties of ASD-associated variants of UBE3C. We selected two documented substitutions: S845F and F996C, respectively localizing to the N- and C-lobe of the catalytic HECT domain (Fig. 2a), a partial structure of which has been resolved ^44^. Both residues are conserved between mouse and human, with only eight amino acids differentiating the entire mouse and human HECT domain of UBE3C.

**Fig. 2.**
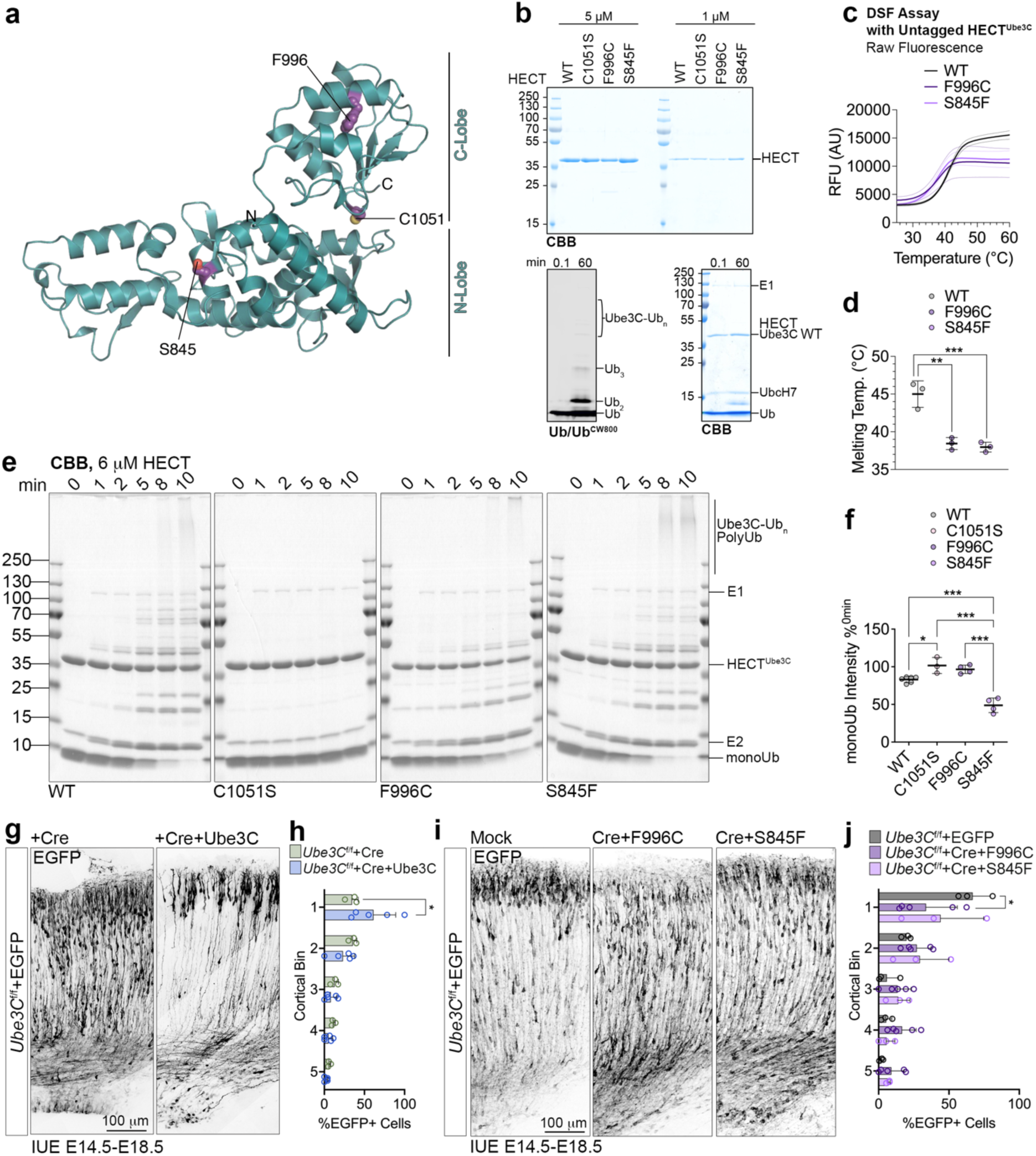
Autism Spectrum Disorder-associated point substitutions in Ube3C alter its stability and enzymatic activity and fail to restore proper cortical assembly. (a) Structure of the HECT domain of UBE3C in an open, L-shaped conformation with its N- and C-lobes and residues substituted in ASD patients, S845 and F996, as well as catalytic C1051S. Side chains are represented as spheres. Structures were folded in AlphaFold 3. (b) Top panel: Coomassie Brilliant Blue (CBB) visualization of the HECT domains variants of Ube3C, including the ASD mutants and synthetic, inactivating point mutation. Bottom panels: autoubiquitination assay of the wild-type (WT) untagged Ube3C HECT domain (HECT^Ube3C^), 1 μM, with UbcH7 as the E2 enzyme, visualized by Ub-CW800 fluorescent probe and CBB. Ube3C-Ub_n_, polyubiquitin chain conjugated to the Ube3C HECT domain. Raw data from biochemical experiments used for representative figures are attached to this manuscript and available on Figshare data repository. (c, d) Differential Scanning Fluorimetry assay with untagged HECT^Ube3C^ variants. RFU, relative fluorescence unit; AU, arbitrary units. (e) Representative images of autoubiquitination assay of HECT^Ube3C^ variants, 6 μM, terminated after indicated reaction time. Reaction products were resolved by SDS-PAGE and visualized by CBB. (f) Quantification of the free monoubiquitin after 5 min of reaction, normalized to the background signal and 0 min timepoint. (g-j) Genetic replacement analyses in *Ube3C* ^f/f^ mouse line to co-express EGFP, Cre and Ube3C variants as indicated using in utero electroporation at E14.5. Brains were fixed at E18.5, sectioned, and immunolabeled using anti-EGFP antibody. (h) and (j) Quantification of laminar positioning of neuronal somata normalized to the global electroporation efficiency per cortex. Individual data points on (d) and (f) represent independent measurements, horizontal line depicts the average and error bars, S.D. Bar graphs and error bars on (h) and (j) represent average and S.D., and data points represent individual electroporated brains used for quantifications. Number of independent measurements for (c, d), 3; for (f), n WT=6, n C1051S=3, n F996C=4, n S845F=4. Number of brains analyzed for (h), n Cre=3, n Cre+Ube3C=5; for (j), n EGFP=3, n F996C=5, n S845F=3. Statistics for (d) and (f), Shapiro-Wilk normality test and one-way ANOVA with Tukey multiple comparisons test; for (h) and (j), two-way ANOVA with Šidák multiple comparisons test. *** p < 0.001; ** 0.001 < p < 0.01; * 0.01 < p < 0.05.

We first recombinantly purified untagged HECT^Ube3C^ variants: wild-type (WT), C1051S (Fig. S1f), F996C, and S845F, and validated the activity of the WT enzyme (Fig. 2b). We then evaluated the thermal stability of the disease-associated HECT fragments using differential scanning fluorimetry (Fig. 2c) and found that both variants are less stable than the WT (Fig. 2d). This implies that the mutated residues contribute to the structural integrity of the HECT domain. Consistently, Phe996 adopts a central position within the hydrophobic core of the catalytic HECT domain C-lobe and Ser845 makes contacts within the N-terminal lobe, including His858 and Gln957.

Based on this finding, we hypothesized that ASD variants might exhibit altered enzymatic activities. To explore this, we assayed their autoubiquitination activities in a reconstituted assay with ubiquitin, ATP, Mg^2+^, E1, and E2 enzyme (Fig. 2e). The C1051S variant served as an inactive negative control (Fig. S1g). Our analysis revealed reduced activity for the F996C variant HECT compared to the WT (Fig. 2f), in line with Phe996 contributing to the structural integrity of the catalytic C-lobe. This demonstrates that ASD-associated mutations may hinder Ube3C function. Intriguingly, the S845F HECT variant exhibited increased activity as compared to the WT. Ser845 assumed the more exposed position on the N-lobe, without involvement in any known binding sites for ubiquitin or the E2 (Fig. 2a, 2e-2f).

Next, we studied the activity of ASD variants in the developing cortex using *Ube3C* ^f/f^ mouse line. In the first experiment, we co-electroporated E14.5 *Ube3C* ^f/f^ cortical progenitors with a Cre-encoding plasmid and with either mock DNA or a construct to re-express wild-type Ube3C (Fig. 2g). Four days later, at E18.5, we found that restoring Ube3C expression reinstates upper layers, compared to the dispersion of cortical neurons across the CP in Cre-expressing cells (Fig. 2h). We then explored the genetic rescue of the cortical lamination using full-length ASD Ube3C variants (Fig. 2i). Notably, the F996C isoform, with lower catalytic activity (Fig. 2e-2f), failed to re-establish upper cortical layers (Fig. 2j). The S845F variant with enhanced activity performed better than F996C in the genetic complementation in vivo; however, it did not fully restore layering in the cortex (Fig. 2i-2j).

Altogether, this integrated analysis of ASD mutations demonstrates that Ube3C catalytic activity correlates with the assembly of cortical neurons into upper cortical layer during embryonic neurogenesis.

### Hyperactivation of Ube3C in dorsal progenitors leads the formation of astrocytes in the developing cortex

Given a previous report on an ASD patient variant with a potential *UBE3C* copy number duplication ^45^, we sought to determine the consequences of its overexpression (OE) in the murine neocortex.

First, we targeted the upper layer cortical lineage using IUE at E15.5 (Fig. 3a) to transiently express EGFP control and Ube3C in cortical progenitors, normally giving rise to upper layer neurons. At P2, control lineages were mostly composed of neurons (Fig. 3b), whereas in cortices electroporated to overexpress Ube3C, we noticed a premature appearance of Sox9-positive astrocytes ^46^.

**Fig. 3.**
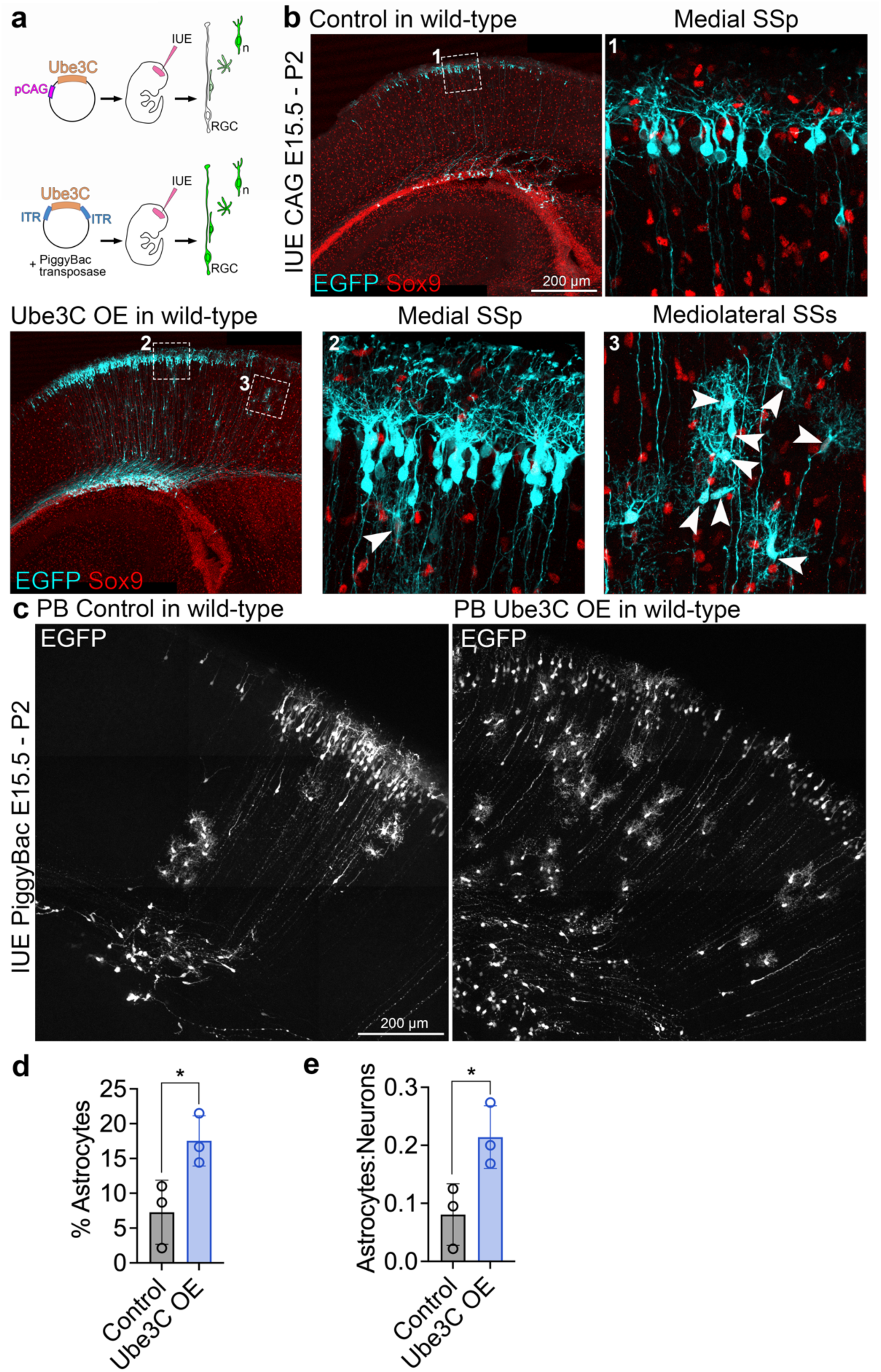
Hyperactivation of Ube3C in cortical progenitors induces precocious astrocytogenesis in the cortex. (a) Schematic depicting the pCAG- and PiggyBac-based in utero electroporation (IUE) experiments to overexpress Ube3C in the developing cortex. IUE using pCAG allows for transient expression of the plasmid in the radial glial cell (RGC) and subsequent inheritance of the plasmid across the RGC lineage, leading to endpoint visualization of neurons (n) in the cortex. PiggyBac system allows for integration of Ube3C into the genome of the entire cortical lineage. (b) EGFP and Sox9 immunostaining in coronal cortical sections from wild-type P2 mice after pCAG-IUE at E15.5 with plasmids to transiently express EGFP and empty plasmid (control) and EGFP and Ube3C (OE) in RGCs. (c) EGFP immunostaining in P2 mice after lineage targeting using piggyBac system at E15.5. White arrowheads in (a) point to EGFP- and Sox9-positive astrocytes. SSp, primary somatosensory area; SSs, supplemental somatosensory area. Note conspicuous astrocytic morphology in spinning disc confocal images. (d, e) Quantification of EGFP-expressing astrocytes in the experiment with piggyBac system, normalized to the global IUE efficiency per cortex (d), or to neuron counts (e). Bar graph and error depict average and S.D., n=3 brains per condition. For statistics, unequal variances t-test. * p < 0.05.

To then stably express Ube3C in the cortical lineage, we leveraged the PiggyBac system (Fig. 3a) ^47^. Contrary to EGFP-expressing controls, among the cells with neuronal morphology, we noticed a significant fraction of astrocyte-like cells in cortices electroporated to overexpress UBE3C (Fig. 3c). Indeed, we then quantified that stable Ube3C OE leads to an increase in the number of astrocytes specified in the cortical progenitor lineage at the expense of neurons (Fig. 3d-3e). Taken together, apart from regulating the laminar distribution of upper layers, Ube3C also controls the identity of cortical progenitor progeny, with loss of Ube3C correlating with neuronal fate (Fig. 1h-1i), and it hyperactivation leading to the appearance of glia (Fig. 3d).

### Ube3C specifies the composition of cortical progenitor lineages in the developing brain

During cortical development, dorsal progenitors first specify neurons and around E16.5 switch to generating astrocytes. Next, we stipulated, that the activity of Ube3C in the cortical progenitors determines the cellular composition of the postmitotic lineage by regulating their neurogenic phase and switch to gliogenesis. To test this, we first explored the molecular landscapes controlled by Ube3C in the developing cortical primordium. The enzymatic activity of Ube3C leads to the formation of a polyubiquitin chain on the target protein. Due to the presence of seven lysine (K) residues on ubiquitin itself, variations in ubiquitin linkages in the polymeric chain dictate the ubiquitin code and determine the fate of the modified substrates ^48^.

For Ube3C, many possible ubiquitin chain conformations have been reported, including branched K29/K48 ^37^, as well as K29-, and K48-linear chains ^49^, the latter canonically associated with protein degradation at the proteasome ^50^. We hypothesized that the loss of *Ube3C* in the developing cortex may lead to a reduction of K48-linked chains assembled on the principal protein targets of Ube3C, and their stabilization.

To explore this possibility, we established Tandem Ubiquitin Binding Entities (TUBEs) affinity chromatography using embryonic cortical tissue, harboring progenitors, as input. TUBEs capture polyubiquitin chains of a given lineage type and allow for the proteomic identification of downstream proteins. We first used Western blotting to demonstrate that K48-TUBEs robustly enriched for K48-polyubiquitinated proteins in cortical lysates from E14.5 control and *Ube3C* cKO (Fig. 4a). We then subjected progenitor-rich E14.5 from control and cKO embryonic cortices to control and K48-TUBEs affinity proteomics (Fig. 4b). Gene Set Enrichment Analysis (GSEA) of the proteins that were significantly less represented in the cKO K48-TUBE sample revealed the regulation of maintenance of sister chromatid cohesion (Fig. 4c), a process critical for the mitotic S phase ^51^. This was a particularly interesting finding, given that cortical progenitors undergo proliferative and consumptive mitoses during neurogenesis to specify their cellular lineages ^52^. We then hypothesized that loss of *Ube3C* might extend the developmental window for consumptive neurogenic divisions of cortical progenitors, diminishing the ability of progenitors to generate astrocytes later in development.

**Fig. 4.**
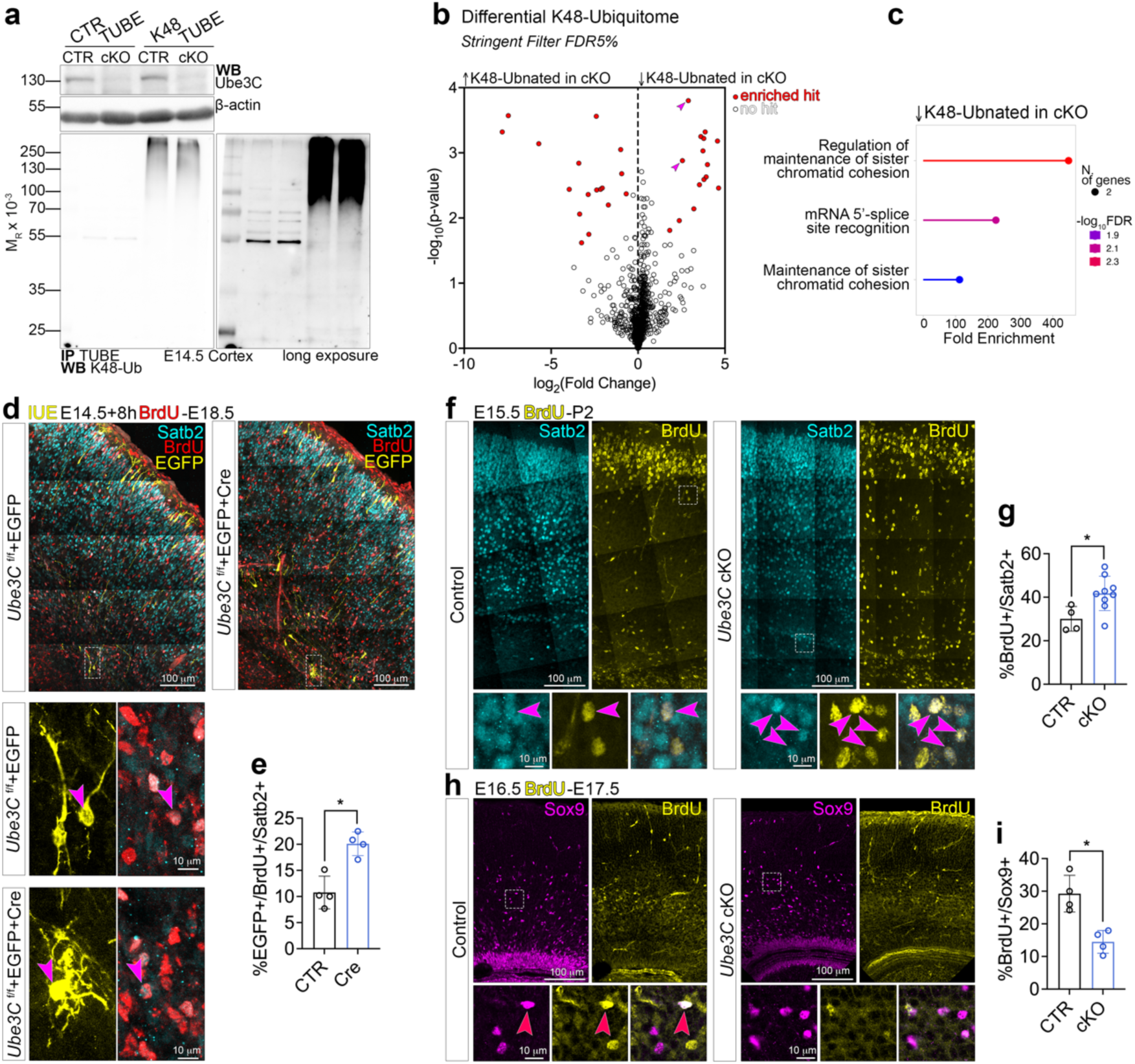
Ube3C supervises the cellular composition of the cortical progenitor lineages by K48-linked ubiquitination of mitotic cycle regulators. (a) Western blotting validation of control- and K48-TUBE-based immunoprecipitation of ubiquitinated proteins in control *Ube3C* ^f/f^ and cKO *Ube3C* ^f/f^; *Emx1*^Cre/+^ E14.5 neocortex. (b) Identification and quantification of K48-ubiquitomes from control and cKO E14.5 neocortex by mass spectrometry. Arrowheads point to regulators of sister chromatid cohesion. (c) Gene Set Enrichment Analysis (https://bioinformatics.sdstate.edu/go/) for proteins identified as enriched hits, with lower K48-ubiquitination mark in *Ube3C* cKO. (d) Representative images of immunostaining signals in E18.5 12 μm-thick coronal cortical cryosections in *Ube3C* ^f/f^ line after in utero electroporation (IUE) with indicated plasmids. BrdU was pulsed intraperitoneally eight hours after IUE. Dotted squares mark insets for higher magnification images. Arrowheads point to triple positive electroporated Satb2+ and BrdU+ neurons. (e) Quantification of triple positive neurons normalized to EGFP+ and Satb2+ double positive cell counts. (f-h) Representative immunostaining signals in 12 μm-thick coronal cortical cryosections of control and cKO mice. BrdU was pulsed at E15.5 (f) or E16.5 (h) and brains were fixed at P2 (f) or E17.5 (h). Arrowheads point to double positive cells and dotted squares represent image insets magnified below. (g, i) Quantification of double positive cells normalized to BrdU+ cell counts. Bar graphs and error bars on (e), (g), and (i) represent averages and S.D., and data points represent individual brains used for quantifications. For (e), n CTR=4, n Cre=4; (g), n CTR=4, n cKO=9; (i), n CTR=4, n cKO=4. For statistics, D’Agostino-Pearson normality test and Mann-Whitney test. ** 0.001 < p < 0.01; * 0.01 < p < 0.05.

Because of this, we explored the cellular fates of progenitors in the *Ube3C* KO mouse line. In the first experiment, we inactivated *Ube3C* using IUE of Cre into E14.5 cortical progenitors in *Ube3C* ^f/f^ mice, followed by intraperitoneal administration of 5’-bromo-2’-deoxyuridine, BrdU, eight hours post-operation (Fig. 4d). BrdU incorporates into the DNA during S phase and allows for birth-dating of specified postmitotic progeny. The vast majority of cells emerging from E14.5 progenitors differentiates into Satb2-expressing neurons, with Satb2 neurons specified across a much broader developmental window ^17,53^. At E18.5, we found that Cre-expressing lineage contained a higher proportion of BrdU-positive, Satb2-positive neurons, indicative of an increase in neurons specified by *Ube3C* KO progenitors (Fig. 4e). We corroborated the increase in neurogenesis in the *Ube3C*-deficient cortical lineages in the *Emx1*-Cre-cKO line as well (Fig. 4f-4g). These results indicate that loss of *Ube3C* enhances neurogenesis and the neuronal output of the cortical progenitors.

During development, upon completion of neurogenesis, radial glial cells (RGCs) switch to producing astrocytes ^7,12^. We hypothesized that enhanced neuron production might deplete the differentiation potential of RGCs later in development, resulting in fewer glial lineages. Notably, we detected fewer Sox9-positive astrocytes specified by cortical progenitors of *Ube3C* cKO mice towards the end of neurogenesis, at E16.5 (Fig. 4h-4i). These findings may indicate that Ube3C regulates the postmitotic cellular diversity of cortical progenitors in part by protracting the neurogenesis window at the expense of gliogenesis.

### Precocious expression of postmitotic transcripts in early *UBE3C* KO organoids

To explore the UBE3C-regulated postmitotic cortical cell fate across species, we then turned towards our organoid model (Fig. S1h-S1j). We first corroborated the viability and development of our organoids at day 30 (30D) using immunolabeling for progenitor identity (Sox9) and *zona occludens* junctions (ZO-1; Fig. 5a). We further corroborated the loss of UBE3C protein from our 30D organoids using Western blotting (Fig. 5b).

**Fig. 5.**
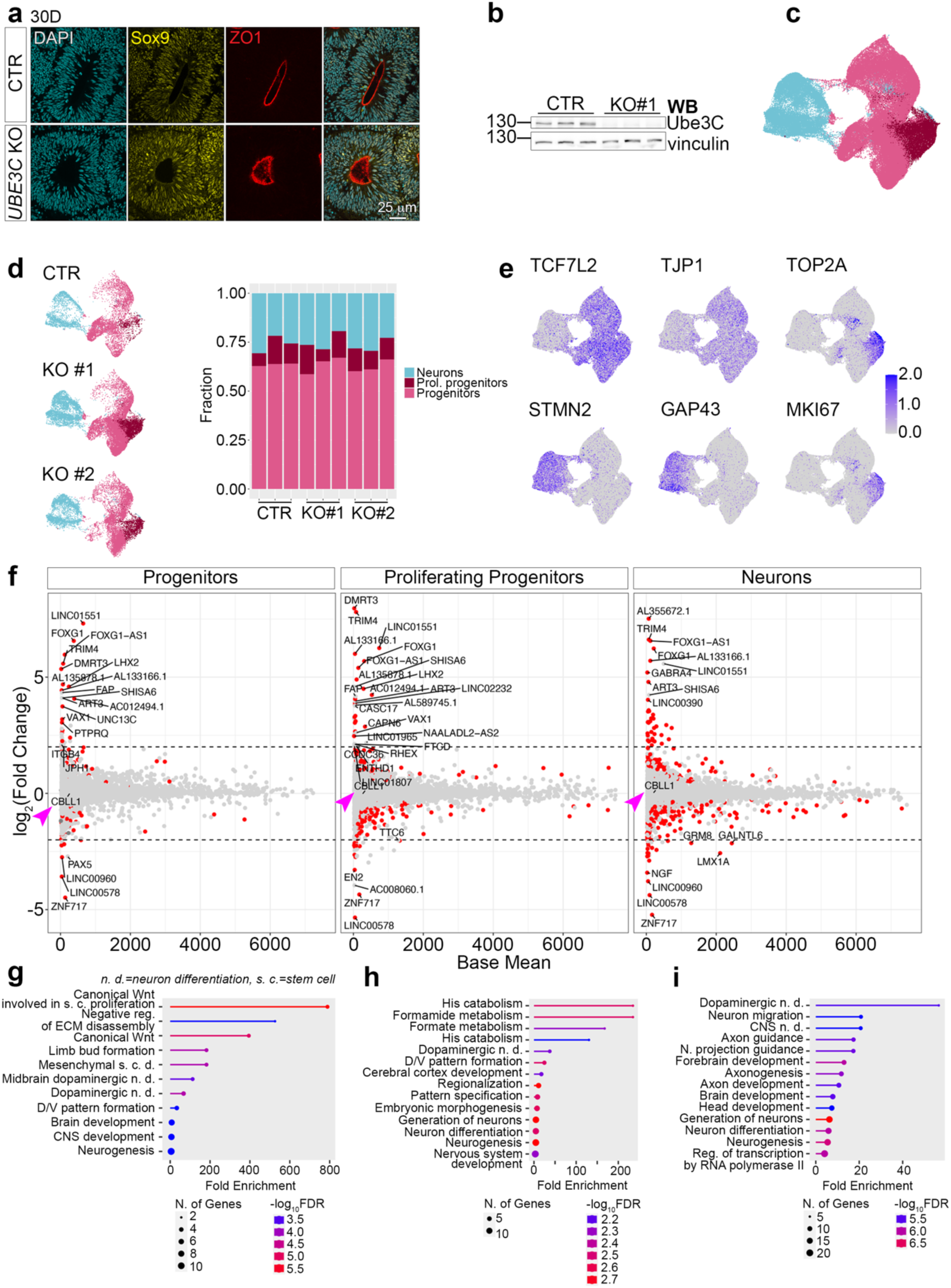
Precocious expression of postmitotic neuron identity in *UBE3C* KO cerebral organoids. (a) Representative images of immunolabeling signals in 30-days-old (30D) control and *UBE3C* KO organoid cryosections. (b) Western blotting validation of *UBE3C* loss in control and KO organoid lysates at 30D. (c) Control (CTR) and KO organoids were subjected to single-nuclei RNA sequencing (snRNAseq). Visualization of identified cell classes using UMAP plot. In total, we analyzed 94 892 single-nuclei with the organoid transcriptomes composed of at least 2000 transcripts collected from three independent replicates, an isogenic control-derived and two independent *UBE3C* KO iPSC-derived organoids. (d) All detected unique cell types were collapsed to three principal cell classes, indicated with different colors. Visualization and quantification of progenitors, proliferating (prol.) progenitors, and neurons in isogenic control (CTR) and *UBE3C* KO organoids derived from two iPSC clones. Each column represents exact values quantified per biological replicate. (e) Expression of mRNA cell identity markers in snRNAseq data. (f) Cell-type specific differential transcript expression in CTR and KO organoids at 30D. Arrowheads point to CBLL1 mRNA. Note unaltered mRNA levels of CBLL1 in KO. (g-i) Gene Set Enrichment Analyses for de-regulated transcripts between CTR and KO for progenitors (g), proliferating progenitors (h), and neurons (i).

To characterize the cellular identities, present in our forebrain organoids, we performed single-nuclei RNA sequencing (sn-RNAseq). We analyzed 94 892 cells, each with at least 2000 transcripts (Fig. 5c), which clustered into three broad cell types: neurons, proliferating progenitors, and progenitors (Fig. 5d), annotated based on the expression of corresponding marker transcripts (Fig. 5e).

Next, within each cell type cluster, we performed differential transcriptome quantification and found increased expression of markers associated with postmitotic neuron fate in *UBE3C* KO organoids, such as FOXG1 and LHX2 ^54^ (Fig. 5f). GSEA in each cell type revealed UBE3C-mediated regulation of neurogenesis (Fig. 5g), differentiation (Fig. 5h), and neuronal migration (Fig. 5i), among others. Altogether, these findings highlight the UBE3C-mediated control of cortical lineage composition also in human tissue, with premature activation of neuronal fate in the KO progenitors.

### Ube3C ubiquitinates Cbll1 to regulate cortical lamination and neuron number

Next, we set out to identify brain-expressed Ube3C targets. The K48-TUBEs approach excludes the ability of Ube3C to assemble K29-linked and other ubiquitin chain types, which are also able to mediate protein degradation ^37,49^. We hypothesized, that the loss of *Ube3C* may lead to a loss of ubiquitination of its principal protein targets, leading to their stabilization and increased expression in the brain. To test this, we used mass spectrometry (MS) to quantify the differential expression of proteins, a method of choice applied in our previous works ^27–29^.

In the MS-based quantification, Ube3C was robustly downregulated, validating its depletion in P0 cortical tissue (Fig. 6a). Among the proteins upregulated in the cKO, representing putative Ube3C ubiquitination substrates, was Cbl Proto-Oncogene Like 1 (CBLL1). Proteins upregulated in the *Ube3C* cKO were associated with developmental maturation, intermediate filaments, cytoskeleton, postsynapse, and mRNA methylation (Fig. 6b). MS-identified hits consist of true Ube3C ubiquitination substrates, predominantly controlled by the ligase, as well as of other proteins, upregulated due to secondary signaling.

**Fig. 6.**
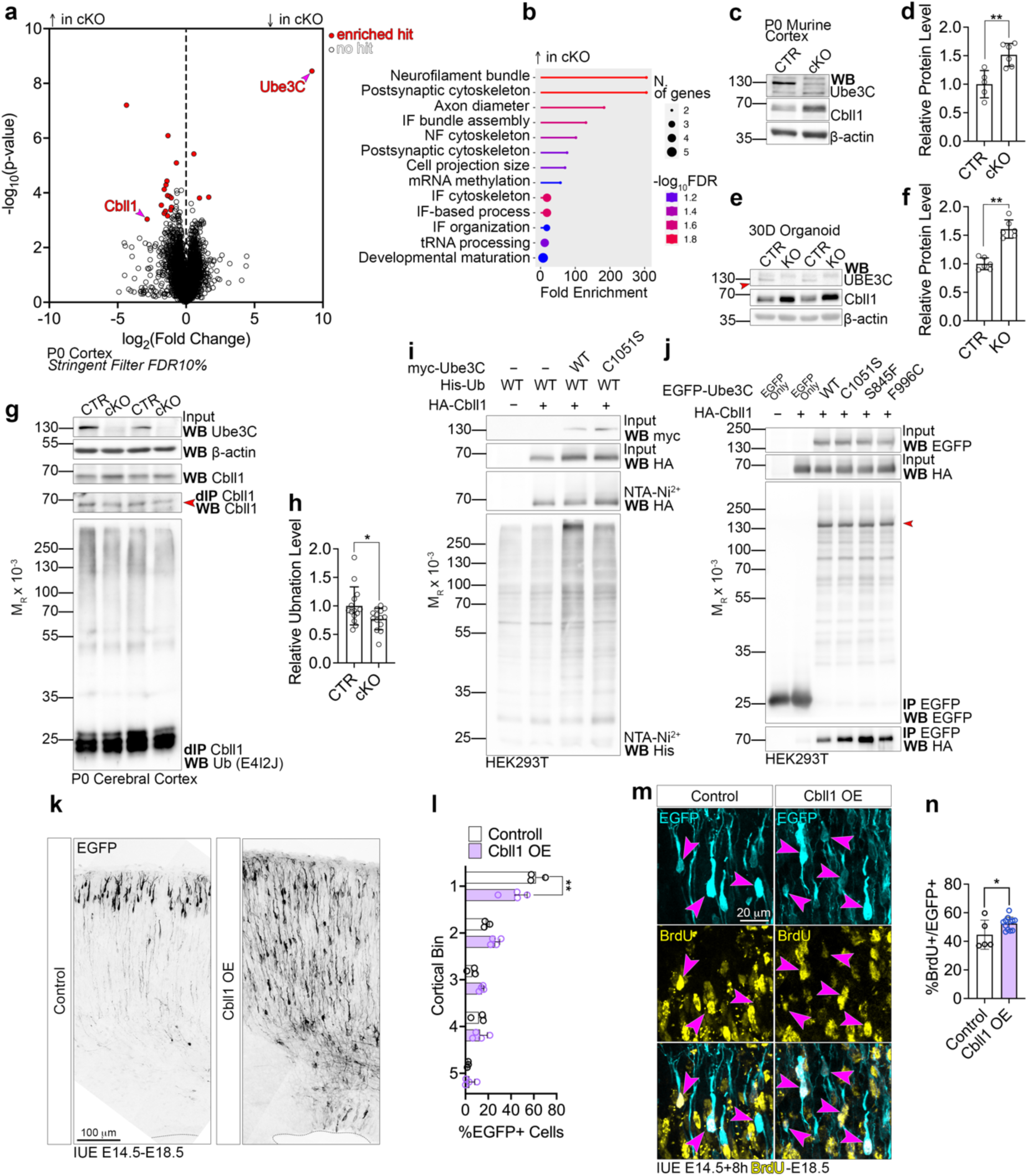
Ube3C-mediated ubiquitination and degradation of Cbll1 regulates the timing of glutamatergic neurogenesis and cortical lamination. (a) Identification and quantification of steady-state proteins in control and cKO neocortex at P0 using Tandem Mass Tag (TMT) mass spectrometry. (b) Gene Set Enrichment Analysis (https://bioinformatics.sdstate.edu/go/) for proteins identified as enriched hits, with higher expression in *Ube3C* cKO cortex. (c-f) Validation of increased Cbll1 expression using Western blotting in P0 murine control and *Ube3C* cKO cortex, as well as 30-day-old control and KO cerebral organoids. Cbll1 level was normalized to beta actin and expressed relative to the control. Arrowhead in (e) points to UBE3C band. (g) Endogenous Cbll1 immunoprecipitation (IP) under denaturing conditions from control and cKO P0 cortex. Note reduced ubiquitination signal in Cbll1 IPed from native cKO cortices. Arrowhead points to Cbll1 band. (h) Quantification of Cbll1 ubiquitination in Western blotting experiment in (g). Ubiquitin signal was normalized to the level of IPed Cbll1. (i) Western blotting results of cell-based ubiquitination assay of transfected HA-tagged Cbll1. (j) Western blotting results of interaction IP between EGFP-tagged Ube3C variants and Cbll1. Arrowhead points to EGFP-Ube3C fusion proteins detected after anti-EGFP IP. (k) Representative images of EGFP fluorescence signals in cortical coronal cryosection of E18.5 wild-type mouse brain after in utero electroporation (IUE) of control plasmids and Cbll1 overexpression at E14.5. (l) Quantification of cortical lamination normalized to the global IUE efficiency in the experiment in (k). (m) Representative images of EGFP and BrdU fluorescence signals in E18.5 neurons electroporated at E14.5 followed by a BrdU pulse eight hours later. (n) Quantification of double positive cells normalized to BrdU+ cell counts. Bar graphs and error bars on (d), (f), (h), (l), and (n) represent averages and S.D., and data points represent individual brains used for quantifications. For (d), n CTR=5, n cKO=6; (f), n CTR=6, n KO=6; (h), n CTR=14, n cKO=14; (l), n CTR=3, n OE=4; (n), n CTR=5, n OE=12. For statistics, (d) and (f), D’Agostino-Pearson normality test and Mann-Whitney test; (h) Anderson-Darling and unequal variances t-test; (l) two-way ANOVA with Šidák multiple comparisons test; (n) Shapiro-Wilk test and unpaired t-test. ** 0.001 < p < 0.01; * 0.01 < p < 0.05.

We next explored Cbll1 as a putative Ube3C substrate. First, we corroborated Cbll1 upregulation in murine P0 *Ube3C* cKO cortical lysate (Fig. 6c and 6d) and human 30D *UBE3C* KO forebrain organoid (Fig. 6e and 6f) by quantitative Western blotting. Importantly, CBLL1 mRNA level was not altered in *UBE3C* KO progenitors or neurons (Fig. 5f, arrowheads), indicating post-transcriptional source of regulation. To test if Cbll1 upregulation was associated with decreased Ube3C-mediated ubiquitination, we established immunoprecipitation (IP) of endogenous Cbll1 from cortical tissue under denaturing conditions. Using an antibody against endogenous protein, we pulled down Cbll1 from control and *Ube3C* cKO P0 cortices (Fig. 6g). We observed a decrease in the polyubiquitin signal normalized to the amount of IPed Cbll1 in the cKO (Fig. 6h). It is important to note that the detected significant difference was observed in vivo, preventing prior stabilization of nascent ubiquitination with DUB inhibitors or proteasome blockers. In the next experiment, we demonstrated that the expression of wild-type Ube3C, but not its catalytically inactive C1051S variant, promotes ubiquitin chain formation on Cbll1 in a cell-based ubiquitination assay (Fig. 6i). Finally, we confirmed a biochemical interaction between Ube3C variants and Cbll1 in a cell-based co-IP experiment. These data indicate that Cbll1 is a brain ubiquitination substrate of Ube3C.

We then explored the physiological relevance of Cbll1 upregulation in the developing cortex. Using IUE, we mimicked the molecular scenario of *Ube3C* depletion by overexpressing Cbll1 in wild-type E14.5 cortical progenitors, and pulsed BrdU eight hours post-operation (Fig. 6k). At E18.5, Cbll1 upregulation induced cortical lamination defects (Fig. 6l), similar to those observed upon Ube3C dysfunction (Fig. 1b-1g). Additionally, we found an increased number of neurons specified in E14.5 progenitor lineages overexpressing Cbll1 (Fig. 6m-6n), resembling the neurogenic phenotype observed in *Ube3C* KO (Fig. 4d-4e).

Altogether, these findings implicate Ube3C-regulated Cbll1 in the developmental control of cortical neurogenesis, influencing both the number of neurons generated by cortical progenitors and their lamination.

### Cbll1 positively regulates the level of cortical m6A mRNA methylation downstream of Ube3C

Next, we explored the molecular mechanisms downstream of Cbll1in the developing brain. Cbll1, also known as Hakai, contains a Really Interesting New Gene (RING) domain and has been described as a ubiquitin ligase controlling E-cadherin endocytosis in epithelial cells ^55^. Intriguingly, in our MS-based quantifications (Fig. 4b and 6a), we did not detect downregulated E-cadherin, or its other described targets, which would be expected for upregulated Cbll1 in the *Ube3C* KO cortex. We therefore hypothesized that in the brain, Cbll1 has other, RING domain-independent, functions.

In fact, Cbll1 has also been shown as a core stabilizer of the N^6^-methyladenosine (m6A) mRNA methylation machinery in *Drosophila*, with Cbll1/Hakai loss leading to impaired m6A installation ^56^. However, the effect of increased Cbll1 levels on m6A writers remained unknown.

Given that the Ube3C-mediated control of cortical cell diversity is embedded in the progenitor cells (Fig. 4), we explored the molecular partners of Cbll1 in the developing cortex at E14.5, when upper layers are specified. We purified protein complexes from the cortical tissue associated with Cbll1 using an anti-Cbll1 antibody and subjected them to MS-based identification (Fig. 7a). The Cbll1 interactome in the developing brain comprised, among others, Vir Like M6A Methyltransferase Associated (Virma) and Cleavage and polyadenylation specificity factor subunit 6 (Cpsf6), molecular factors regulating m6A writers.

**Fig. 7.**
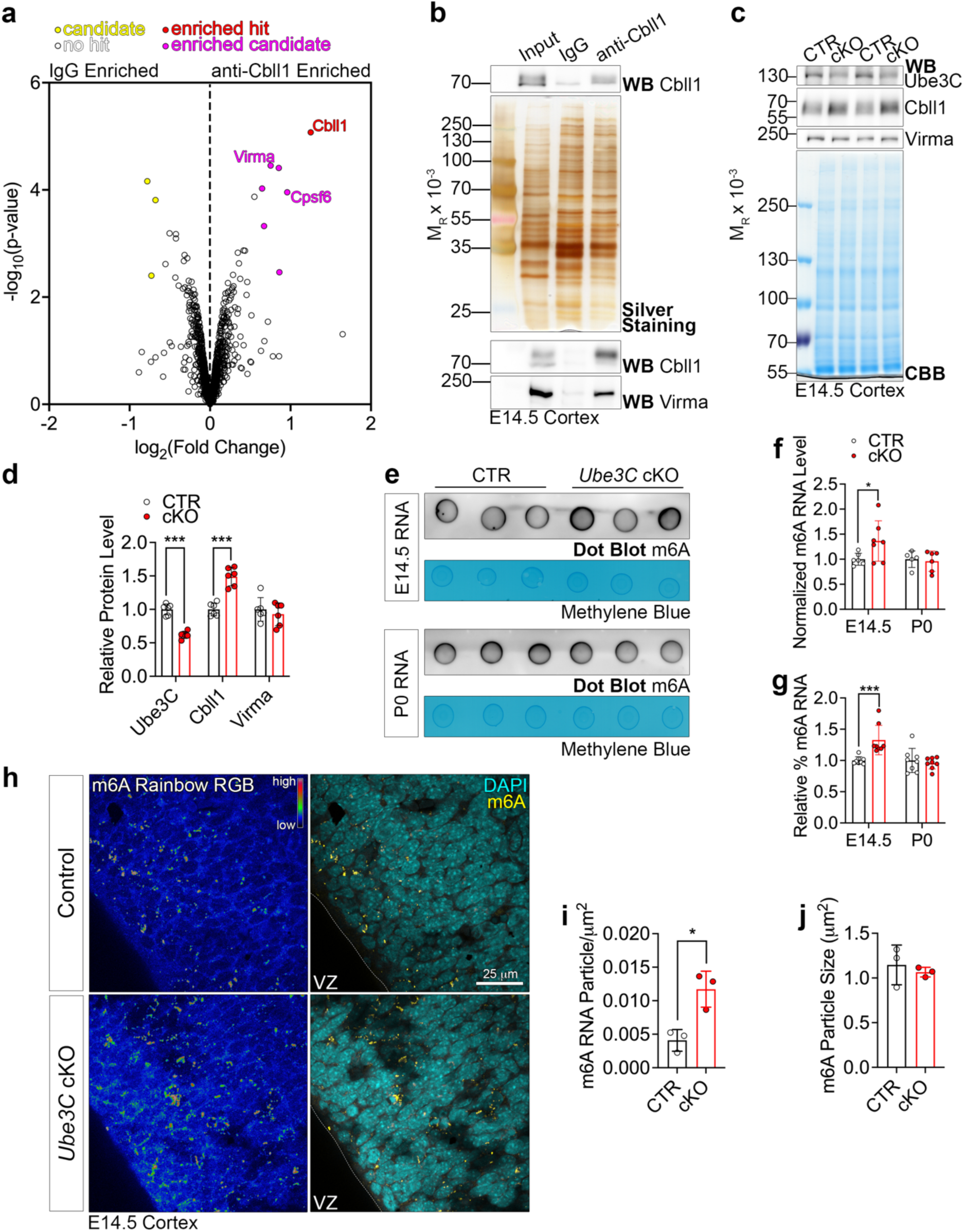
Ube3C-regulated Cbll1 drives the activity of N^6^-methyladenosine (m6A) writers in the embryonic developing cortex. (a) Results of quantitative mass spectrometry to identify protein interactome of Cbll1 in IgG- and anti-Cbll1 immunoprecipitation (IP) from E14.5 wild-type cerebral cortex using mass spectrometry. (b) Validation of the anti-Cbll1 IP using cortical lysate input, as well as validation of interaction between Cbll1 and Virma in the E14.5 cortical lysate. (c) Representative Western blotting results for the levels of indicated proteins in E14.5 control, CTR, and *Ube3C* cKO cortical lysates. CBB, Coomassie Brilliant Blue. (d) Quantification of protein levels normalized to CBB and expressed as fraction of control. (e) Representative Dot Blot experiment to detect m6A in RNA isolated from CTR or cKO at indicated developmental timepoints. (f) Quantification of m6A signal normalized to the Methylene Blue stain and expressed as a fraction of control per developmental stage. (g) Quantification of m6A content in total RNA isolated from CTR and cKO cortex using fluorometric assay. (h-j) Representative fluorescence signals from m6A (represented as RBG Spectrum) and DAPI from CTR and cKO E14.5 cortex and quantification of particle density (i) and size (j). VZ, ventricular zone. Bar graphs and error bars on (d), (f), (g), (i), and (j) represent averages and S.D., and data points represent individual brains used for quantifications. For (d), n CTR=6, n cKO=6; (f), E14.5, n CTR=6, n cKO=7; P0, n CTR=5, n cKO=6; (g), E14.5, n CTR=8, n cKO=8; P0, n CTR=8, n cKO=8; (i), n CTR=3, n cKO =3; (j), n CTR=3, n cKO =3. For statistics, (d), Shapiro-Wilk test and unequal variances t-test; (f), D’Agostino-Pearson normality test and unpaired Kolomogorov Smirnov test; (g) D’Agostino-Pearson normality test and Mann-Whitney test (E14.5) or Welch’s t-test (P0); (i) and (j), Shapiro-Wilk test and Welch’s t-test. *** p < 0.001; * 0.01 < p < 0.05.

Virma functions as a scaffold protein within the METTL Associated Complex (MACOM), essential for the full catalytic activity of the METTL3/METTL14 heterodimer^57^. METTL3 catalyzes the co-transcriptional deposition of m6A on mRNA. Cpsf6, a poly-adenlyation factor, is functionally linked to Virma and their interaction facilitates the specific deposition of m6A on mRNAs ^58^. We hypothesized that increased Cbll1 in *Ube3C* cKO might result in enhanced activity of m6A writers.

To explore this possibility, we first corroborated the interaction between Cbll1 and Virma in the E14.5 embryonic cortex using co-IP experiment (Fig. 7b). Cbll1 levels were upregulated in E14.5 *Ube3C* cKO cortex, shown by quantitative Western blotting, unlike Virma protein expression, which remained unaltered (Fig. 7c-7d). Next, we employed a dot blot assay to quantify the level of m6A modification in RNA samples, purified from control and *Ube3C* cKO cortex at E14.5 and P0 (Fig. 7e). Intriguingly, we found increased m6A levels in cKO RNA sample at E14.5, but not at P0 (Fig. 7f). We then used a colorimetric assay to quantify the proportion of m6A within the total RNA sample and corroborated previous results of the dot blot experiment results (Fig. 7g). Finally, we used an immunohistochemistry-based approach to visualize m6A in cortical cryosections of control and *Ube3C* cKO brains (Fig. 7h). Consistently, we found increased density of m6A particles in the cKO (Fig. 7i), with no alternations in the size of the m6A deposits (Fig. 7j). These results indicate that Ube3C-Cbll1 axis drives m6A deposition on RNA specifically during cortical neurogenesis at E14.5.

### Pharmacological inhibition of METTL3 restores cortical neurogenesis in *Ube3C* KO mice

Having understood the principal molecular pathway regulated by Ube3C-Cbll1 axis, we explored a possibility of rescuing the aberrances in cortical development associated with *Ube3C* depletion. Preclinical studies have shown that MAC complex inhibition via a small molecule STM2457 suppresses the growth of neuroblastoma tumors in vivo ^59^. We hypothesized that increased neuron number specified by *Ube3C* KO (Fig. 4d-4e), cKO (Fig. 4f-4g), and Cbll1 OE (Fig. 6m-6n) progenitors are attributable to the aberrances in the cell cycle exit during neurogenesis. To test this, we intraperitoneally administered a single dose of 50 mg/kg STM2457 in DMSO ^59^ to pregnant E14.5 *Ube3C* ^f/f^ mother mated to *Ube3C* ^f/f^ male. We used IUE to electroporate cortical progenitors with either EGFP (control), or to co-express EGFP and Cre (KO). At E15.5, we determined the pool of cells which exited the mitotic cycle, using immunohistochemistry for Ki67 proliferation marker (Fig. 8a). In vehicle-exposed *Ube3C* KO brains, we found increased fraction of cycling progenitors, as compared to DMSO-exposed control ones, which might represent the increased neurogenic potential of *Ube3C* KO neural stem cells. Strikingly, exposure of *Ube3C* KO progenitors to STM2457 restored the postmitotic specification in the cortical lineage (Fig. 8b). This result supports the potential of METTL3 inhibition in restoration of NDD-associated developmental aberrances in *Ube3C* KO mouse model.

**Fig. 8.**
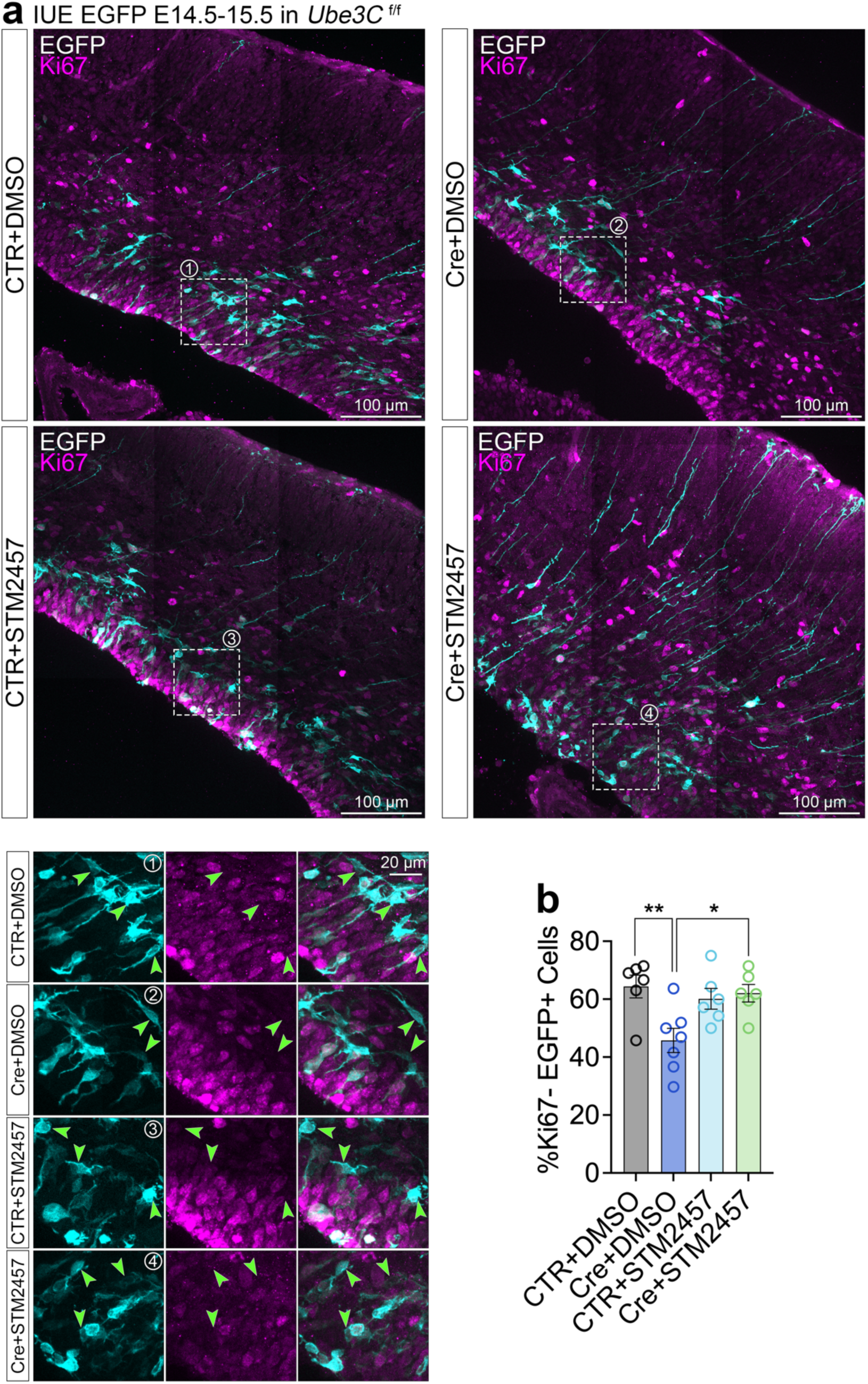
Inhibiting m6A writing in *Ube3C*-deficient cortical progenitors restores aberrant dynamics of cell cycle exit during neurogenesis. (a) Representative fluorescence signals of EGFP and Ki67 in E15.5 cortical coronal cryosections from *Ube3C* ^f/f^ mouse brains electroporated with EGFP or EGFP and Cre, and DMSO- or STM2457-pulsed eight hours post-surgery. Dotted squares indicated magnified regions represented in the insets below. Arrowheads point to postmitotic EGFP+ cells. (b) Quantification of the proportion of postmitotic EGFP+ cells normalized to the EGFP-positive cell number per section per brain. Bar graphs and error bars on (b) represent averages and S.D., and data points represent individual brains used for quantifications. For (b), n CTR+DMSO=6, n Cre+DMSO=7, n CTR+STM2457=6, n Cre+STM2457=6. For statistics, Kolomogorov Smirnov test and one-way ANOVA with Tukey’s multiple comparisons test. ** 0.001 < p < 0.01; * 0.01 < p < 0.05.

## DISCUSSION

Neuronal development is a tightly coordinated process with cellular transitions relying on high-order signaling pathways to ensure swift remodeling of molecular landscapes in transitory cellular niches. E3 ubiquitin ligases, as substrate specificity determinants, regulate the spatiotemporal control of ubiquitination, altering cellular protein levels through proteasomal, autophagosomal, or lysosomal degradation ^11^. We are only beginning to appreciate the vast biological processes governed by ubiquitination and E3 ligases, which, alongside extensive studies on protein synthesis, have emerged as a distinct subfield in neurodevelopment ^26^. Notably, nearly one-eighth of all known E3 ligases are directly linked to severe genetic neurological disease ^60^. The urgency of understanding the basic E3 ligase function is underscored by the fact that neurological conditions are the leading cause of illness and disability, affecting over one-third of the global population ^61^.

We discovered that the HECT ligase Ube3C dictates cortical cell types generated by neural stem cells (Fig. 3 and 4). In our *Ube3C* cKO mouse model and in IUE-driven single lineage depletions, excess neurons fail to organize into the characteristic single-layer structure of the mammalian cerebral cortex (Fig. 1, 2, and 4). This is particularly significant, as UBE3C mutations have been identified in NDD patients with varying severity and symptoms. Loss of *UBE3C* leads to severe NDD with ID ^34^, while point mutations in the catalytic domain are linked to sporadic ASD ^35,36^. Through integrated analyses combining analytical biochemistry and in vivo genetics, we report severe neurodevelopmental aberrations for complete *Ube3C* loss (Fig. 1) and the F996C variant, which exhibits reduced autoubiquitination activity (Fig. 2). These results align with emerging evidence that molecular changes in neural stem cells correlate with circuit-level defects in the adult NDD brain ^6^. Thus, Ube3C - and recently also Ube3A^62^ - links NDDs, including ASD, to a progressive multistage disorder where prenatal neurodevelopmental defects disrupt cortical lamination and cellular diversity, ultimately affecting cortical network assembly in adulthood. The distinct developmental effects of the F996C and S845F mutations may contribute to the biological basic of syndromic ASD heterogeneity.

Our enzymatic assay used purified HECT domains, as full-length Ube3C generation is technically challenging (Fig. 2a-2g). Our findings suggest that F996C in the C-lobe of the HECT domain affects the conformational flexibility required for ubiquitin conjugation (Fig. 2a and 2e-2g). Although the S845F mutation seems to be redundant for cortical lamination, it exhibits decreased thermal stability (Fig. 2d). We speculate that this mutation in physiological context, albeit increased in vitro activity, may lead to reduced stability.

We describe a dual developmental role for Ube3C, promoting neurogenesis at the expense of astrogenesis (Fig. 3 and 4), resulting in excess neurons, which fail to laminate and form layer II/III (Fig. 1 and 4f). Given that the targeted E14.5 cortical lineage differentiates predominantly to Satb2-positive neurons ^17^, the increased neuronal production in *UBE3C*-depleted models may represent a protracted neuron production window ^53,63^. This is further supported by precocious induction of pro-neuronal fate transcripts in early *UBE3C* KO organoid (Fig. 5). As progenitor pools become gradually exhausted, the sustained cell cycle activity may result in reduced gliogenesis. At the molecular level, transcriptomic changes in forebrain organoids converge on the regulation of neuronal migration (Fig. 5i), while proteomic analyses implicate Ube3C in cytoskeleton regulation (Fig. 6b), suggesting that disrupted radial migration of neurons is the key cause of cortical lamination defects, the second developmental milestone regulated by Ube3C.

Through integrated proteomics, we discovered Cbll1 as the principal Ube3C target in the developing brain. Our experimental design constrained proteomic identification to substrates predominantly controlled by Ube3C (Fig. 4b and 6a). However, Cbll1 overexpression in wild-type cortical progenitors replicated the developmental defects observed in *Ube3C* KO mice (Fig. 6k-6n). Furthermore, linking m6A deposition to Cbll1 signaling (Fig. 7) enabled pharmacological restoration of Cbll1-driven *Ube3C* KO abnormalities (Fig. 8). Increased Cbll1 levels in *Ube3C* KO implicate its degradation (Fig. 6a-6f, 7c-7d), likely at the proteasome, although we did not detect Cbll1 in the TUBE experiment (Fig. 4b), hinting that Ube3C conjugates non-K48-linked polyubiquitin chains on Cbll1. Ube3C has been shown to catalyze K29-linked chains, associated with degradation in other tissues ^37,49^. Additionally, Ube3C regulates proteasome activity independently of its canonical E3 function ^64^, further highlighting the complexities of molecular dependencies in the brain.

Cbll1 has been reported as a RING-type E3 ubiquitin ligase in epithelial tissue ^55^. We speculate that upstream regulators and molecular interactions may render its ligase activity in a tissue-defined context. Our mass spectrometry analysis of Cbll1 interactors identified Virma and Cpsf6 (Fig. 7a-7d), components of MACOM complex. Given that MACOM is disrupted upon Cbll1 loss ^56^, it is plausible that increased m6A seen in our model is linked to hyperstabilization of the MACOM complex and enhanced METTL3 enzymatic activity, when Cbll1 is overexpressed (Fig. 6c-6d, 7c-7d). Intriguingly, we found elevated m6A deposition on RNA specifically at the embryonic stage (Fig. 7e-7j). It is conceivable that this represents the activity of the Ube3C-Cbll1axis in E14.5 cortical progenitors. As is the case for other HECTs, Ube3C undergoes post-translational modification-driven spatiotemporal regulation ^38^. Postnatally, Ube3C-Cbll1 duo may govern distinct processes, such as cytoskeleton dynamics, as indicated by GSEA for upregulated proteome in *Ube3C* cKO (Fig. 6b). Future follow-up studies should characterize the m6A-regulated transcriptome in *Ube3C* KO models.

Finally, our discovery that a single dose of STM2457 to inhibit METTL3 restores progenitor cell cycle dynamics in *Ube3C* KO lineages presents a potential therapeutic avenue (Fig. 8). As a multidisciplinary pipeline, our findings provide a comprehensive understanding of Ube3C-regulated signaling and may pave the way for genetic defect-targeted interventions ^32^. However, potential postnatal therapeutic strategies remain unexplored, as the Ube3C-Cbll1 axis primarily functions in the embryonic cortex to modulate m6A installation on mRNA (Fig. 7).

Alongside transcription, mRNA splicing, protein translation and stability regulation, E3 ligase-mediated ubiquitination and degradation have emerged as key regulators of gene expression in neurodevelopment. In the context of cellular composition of the brain and regulation of developmental tempo ^63,65^, the E3s represent a widespread theme, given their extensive substrate repertoire and signaling influence. In the foreseeable future, integrating diverse scientific subfields will illuminate how interconnected fundamental processes - such as ubiquitination, ubiquitin-like modifiers, mitophagy, metabolism, vesicle sorting, and proteins biosynthesis - coordinate cellular mechanisms organizing neurodevelopmental transitions ^32,66–68^.

## ACKNOWLEDGEMENTS

We thank Fritz Benseler, Jutta Schüler, Claudia Abramjuk, Raimunde Hellwig-Träger, and Dietmar Schmitz for their support and help with this project. We acknowledge the support by the BIH Cytometry Core Facility and the Advanced Medical Bioimaging Core Facility of the Charité. We thank Andrew Newman for his helpful tips on the experiments with m6A, Georgia Rapti and Marta de Rocha Rosário for their conceptual advice. This project was funded by the German Research Foundation, Deutsche Forschungsgemeinschaft DFG, project number 536563141 (to MCA). MCA is a scholar of the FENS-Kavli Network of Excellence and work in his lab is supported by the Fritz Thyssen Foundation (10.23.2.003MN) and the DFG (515247130). PM was supported by the DFG, under the grant agreement SFB 1588, project A06, "Neuroblastoma Evolution". DS was supported by the DFG Emmy Noether Programme (SCHW1851/1-1). SL acknowledges support from the Max Planck Institute for Multidisciplinary Sciences, the Max Planck Society, and the SFB1565 (DFG; 469281184, P17). HK was supported by the Takeda Science Foundation.

## AUTHOR CONTRIBUTIONS

MCA conceptualized the project, acquired and analyzed data, compiled figures and wrote the manuscript with input from other authors. KB carried out the bulk of IUE experiments, histological characterizations, acquired and analyzed microscopy data. KJC characterized the ASD UBE3C mutants, performed the BrdU experiments and investigated Ube3C-CBll1 functional interaction in the mouse and organoids. RD performed most of biochemical experiments in this manuscript, as well as m6A assays. JK carried out the autoubiquitination assays of Ube3C variants. TS performed most of the molecular cloning for the plasmids generated in this study. JN and MS helped with data acquisition and analysis. Organoid work was supervised and partially carried out by ARW. NK generated *UBE3C* KO iPSC lines and performed quality check and organoid imaging experiments. IK helped with organoid derivation, initial characterization and immunostaining. TMT proteomics and TUBE mass spectrometry was performed by MK and PM. Cbll1 interactomics was carried out by FS and PH. RNA sequencing was performed by the Genomics Core Facility, CD, CQ, TB, and TC. TP and NR analyzed snRNAseq data. DS helped with the initial purification of GST-tagged UBE3C variants. TF and SL purified untagged UBE3C variants and carried out the DSF experiments. HK and NB initialized the project and supervised the Ube3C mouse line generation. VT enabled and supported the experimental work and supervised the work in his lab.

## ETHICS DECLARATION

The authors declare no competing interests.

## MATERIALS AND METHODS

### Animals

Mouse (*Mus musculus*) lines described in this study were maintained in the animal facilities of Max Planck Institute for Multidisciplinary Sciences (MPI-NAT) and the Charité. Mice were housed in a 12 h light/dark cycle at 18-23°C, 40-60% air humidity, with pellet food and water available *ad libitum*. Mice were used at embryonic (E), or postnatal (P) stages, as reported for each experiment. *Ube3C* ^f/f^ mouse line was generated in this study by homologous recombination in ES cells (The European Conditional Mouse Mutagenesis Program, EUCOMM). Animals homozygous for the *loxP* alleles were viable, fertile, born at the expected Mendelian ratio, and exhibited no overt phenotypic changes in the cage environment. To inactivate *Ube3C* in the developing cortex, we crossed *Ube3C* ^f/f^ mice with the *Emx1*-Cre line (B6.129S2-Emx1^tm1(cre)Krj^/J), in which Cre recombinase is expressed from the *Emx1* gene allele^40^. *Ube3C* ^f/f^ and *Ube3C* ^f/f^; *Emx1*^Cre/+^ were maintained in C57BL/6N background. Wild-type mice were of NMRI strain. Females were housed in groups of up to five animals per cage. Males were single housed from the point of exposure to female animals. The date of vaginal plug was counted as E0.5. All mice were sacrificed by administering a lethal dose of pentobarbital.

### Inclusion and ethics statement

MCA is a signatory of the ALBA Declaration on Equity and Inclusion. All experiments were performed in compliance with the guidelines for the welfare of experimental animals approved by the State Office for Health and Social Affairs, Council in Berlin, Landesamt für Gesundheit und Soziales (LaGeSo), permissions: G0079/11, G0206/16, G0184/20, G0054/19, G0055/19, G0019/24, T102/11, and T-CH-0033/22.

### Sex and age/developmental stage of experimental animals

Littermates of both sexes were chosen for the biochemical and histological analyses, according to the SRY allele genotyping ^69^. Each sample set contained an even number of male and female samples. Developmental stages or stages at experimental interventions are listed on the figures or in the figure legends.

### Mouse Embryonic Fibroblasts (MEFs)

Neomycin-resistant MEFs (Cell Biolabs, CBA-311) were used as feeder cells for embryonic stem (ES) cell culture. MEFs were cultured in 15 cm Petri dishes coated with gelatin (0.1% gelatin/ddH_2_O) at 37°C in 25 mL MEF medium in the presence of 5% carbon dioxide in HERA-cell240 (Heraeus) incubator. At confluency, cells were washed with 37°C-warm PBS, and incubated with 10 mL MEF medium supplemented with 100 μL of mitomycin C (Sigma-Aldrich) per plate for 2.5 hours at 37°C to mitotically inactivate MEFs. Then, medium was removed, cell washed with PBS, trypsinized for 5 minutes with 0.05% trypsin-EDTA solution (Gibco, Life Technologies) at 37°C, resuspended with fresh MEF medium, and centrifuged at 800 g for 5 minutes at room temperature. Inactivated MEF were then frozen in freezing medium and Cryo Freezing Container (Nalgene) filled with isopropanol at -80°C, and seeded on demand.

MEF medium: 500 mL KnockOut-DMEM (Gibco, Life Technologies), 75 mL FBS, 6 mL non essential amino acids (Gibco, Life Technologies), 6 mL 200 mM L-glutamine (Gibco, Life Technologies), 6 mL β-mercaptoethanol (Sigma-Aldrich), 3 mL penicillin/streptomycin, 6 mL G418 (Gibco, Life Technologies).

Freezing Medium: 22 mL MEF medium, 10 ml FBS, 8 mL dimethyl sulfoxide (DMSO, Sigma-Aldrich).

### Embryonic Stem Cells (ESCs)

For generation of *Ube3C* knock-out mice, mutant JM8A3.N1 ES cells were purchased from EUCOMM Consortium (Ube3C^tm1a(EUCOMMHmgu)^). The L1L2_Bact_P cassette was inserted upstream of exon 5 of *Ube3C* gene. The cassette was composed of an FRT-flanked lacZ/neomycin sequence followed by a 5’ loxP site. An additional *loxP* site is inserted 3’ downstream of exon 5. ES cells were thawn prior to blastocyst injection to minimize the passage number. Cells were seeded on a layer of previously prepared and thawn inactivated MEF, and maintained in ES cell medium at 37°C in the presence of 5% carbon dioxide in HERA-cell240 (Heraeus) incubator. Before splitting, cells were supplied with fresh medium for exactly 3 hours. Passaging was carried out with 0.25% trypsin-EDTA (Gibco, Life Technologies), and the reaction was terminated with FBS. Prior to blastocyst injection, MEFs were separated from ES cells by plating cell suspension on fresh gelatinized Petri dish for 30 minutes at 37°C allowing MEFs to adhere to the bottom of the dish. Transfer of injected blastocysts to pseudo-pregnant foster female mice was performed in MPI-NAT Animal House Facility.

ES cell medium: 500 mL KnockOut-DMEM, 95 mL FBS, 6 mL non essential amino acids, 6 mL 200 mM L-glutamine, 6 mL β-mercaptoethanol, 3 mL penicillin/streptomycin, 6 mL G418, 1000 U Leukemia Inhibitory Factor (Chemicon/Millipore).

### Generation of *Ube3C* ^f/f^ mouse line

ESCs were validated with PCRs and southern blotting (Fig. S1b-S1e) for the correct insertion of the targeting cassette and injected into recipient blastocysts. Mice derived from such ES cells were crossed with flip recombinase expressing mouse line (B6;SJL-^Tg(ACTFLPe)9205Dym^/J; Jackson Laboratory) to delete the lacZ/neomycin cassette. *Ube3C* ^f/+^ mice were crossed to establish *Ube3C* ^f/f^ mouse line.

### Cell lines

HEK293T cells lines were from the Leibniz Institute DSMZ (https://www.dsmz.de/) and were maintained in the standard medium. The cells were routinely tested negative for mycoplasma contamination prior to the experiments.

<u>HEK cell medium</u>: 500 mL DMEM, 50 mL FBS, 5 mL GlutaMAX, 5 mL penicillin/streptomycin (from 100X). All components were from Gibco, Life Technologies.

### PCR reaction (Ube3C ^f/f^ mouse line genotyping, diagnostic PCRs)

PCR for Ube3C-*flox* allele was carried out using oligonucleotides listed below and the following parameters. Reaction with *flox*-allele yields a band of 338 base pairs (bp); wild-type allele – 274 bp.

Primer Ube3_P1: 5’-CTCACAGGGCAGTAATTGTCTC-3’

Primer Ube3_P2: 5’-TAATCTGAAGTTGCCTAAACTCCT-3’ Primer Ube3_P3: 5’-CCACAACGGGTTCTTCTGTTAG-3’

Step 1: 96°C for 3 minutes,

Step 2: 94°C for 30 seconds,

Step 3: 64°C for 60 seconds,

Step 4: 72°C for 60 seconds (33 cycles from Step 2 to 4)

Step 5: 72°C for 7 minutes.

Multiplex PCR to detect 3’loxP site (Fig. S1d) was performed with these parameters as well. Amplification of long genomic DNA sequences for validation of correct recombination of targeting vector to genomic DNA, (Fig. S1c) was performed using Pfu-AD polymerase in Phusion-HF buffer (NEB) in the presence of 1 M betaine (Sigma-Aldrich) with the following parameters:

Step 1: 99°C for 3 minutes,

Step 2: 99°C for 30 seconds,

Step 3: 60°C for 30 seconds,

Step 4: 72°C for 90 seconds (30 cycles from Step 2 to 4)

Step 5: 72°C for 2 minutes.

### Southern blotting

The procedure for southern blot in this paper was exactly as described in the Supplementary material in ^70^. Radioactive probes were synthetized with Prime-It II Random Primer Labeling Kit (Agilent) using 25 ng of EcoRI-digested template and α-^32^P-labeled dCTP (Perkin-Elmer) following manufacturer’s protocol. These experiments were carried out by MS.

### Molecular cloning strategies for constructs generated in this study

Cloning of most expression vectors in this study was performed using the NEBuilder system according to the manufacturer’s protocol (New England BioLabs). DNA fragments were amplified using GXL Prime Star DNA polymerase (Takara) using cDNA libraries or donor plasmids as templates, as indicated.

#### pCR2.1-Ube3C (WT, full length)

cDNA NM_133907.3 encoding full length (1083 amino acids) murine Ube3C (Origene, SKU MR223355L4) was amplified with primers: 5’-GAGGAATTCACCATGTTCAGCTTCGAAGGCGACT-3’ and 5’-CTCGTCGACTTATCAGCTCAGCTCAAAGCCAGC-3’, introducing 5’EcoRI and 3’SalI restriction sites using pLenti-Ube3C as a template (Origene, SKU MR223355L4). Resulting DNA was then subcloned into pCR2.1-TOPO.

#### pCR2.1-SB-5’Ube3C

To generate Southern blot probe for validation of the correct recombination of Ube3C targeting vector, the following primers were used for PCR using DNA isolated from wild-type ES cells as a template: 5’-TTGTGAGCTTTTCCTTTTGTCCA-3’ and 5’-TTTAGAGAAATGGGTTATAATGTTTTAA-3’. Amplified DNA was subcloned into pCR2.1-TOPO.

#### pCR2.1-SB-3’Ube3C

To generate Southern blot probe for validation of the correct recombination of Ube3C targeting vector, the following primers were used for PCR using DNA isolated from wild-type ES cells as a template: 5’-GACCATGATGCAAGATGTGG-3’ and 5’-CTTGACTAGTATATCCTCACTGCTAGTT-3’. Amplified DNA was subcloned into pCR2.1-TOPO.

#### pCAG-myc-Ube3C

Upon EcoRI/SalI restriction digest of pCR2.1-Ube3C, the purified resulting insert was ligated to pCAG-myc (pRaichu-myc), opened with EcoRI and XhoI.

#### pCR2.1-Ube3C C1051S

The point mutation to replace the catalytic cysteine to serine (3151T>A) was introduced using the Site-Directed QuickChange Mutagenesis Kit (200523, Agilent) and the following primers: 5’-TTGAGCGGCTACCTACAGCCAGCACCAGCATGAACCTGC-3’ and 5’-GCAGGTTCATGCTGGTGCTGGCTGTAGGTAGCCGCTCAA-3’.

#### pCAG-myc-Ube3C C1051S

Upon EcoRI/SalI restriction digest of pCR2.1-Ube3C C1051S, the purified resulting insert was ligated to pCAG-myc (pRaichu-myc), opened with EcoRI and XhoI.

#### pCAG-myc-Ube3C S845F

The mutant was created from a plasmid template pRai-myc-Ube3C using the following oligos: 5’-gtctcatcattttggcaaagATGGAACAAAAACTCATCTCAG-3’ with 5’-ggaaaaagccAGCAAAGGGTAGCTCCAC-3’ and 5’-accctttgctGGCTTTTTCCTGTTCAAGC-3’ with 5’-cggccgcgatatcctcgaggTCAGCTCAGCTCAAAGCCAGCA-3’ and inserted into the pCAG-expression vector (Addgene, #11160) using EcoRI restriction and NEBuilder system (E2621L, NEB).

#### pCAG-myc-Ube3C F996C

The mutant was created from a plasmid template pRai-myc-Ube3C using the following oligos: 5’-gtctcatcattttggcaaagATGGAACAAAAACTCATCTC-3’ with 5’-ttttaatgacAGGATGATCGGCAGAATAG-3’ and 5’-cgatcatcctGTCATTAAAATCTTTTGGAGAGTTGTAG-3’ with 5’-cggccgcgatatcctcgaggTCAGCTCAGCTCAAAGCCAGCATCAGCTCAGCTCAAAGC C-3’ and inserted into the pCAG-expression vector (Addgene, #11160) using EcoRI restriction and NEBuilder system.

#### pET-Ube3C-HECT WT, pET-Ube3C-HECT C1051S, pET-Ube3C-HECT S845F, pET-Ube3C-HECT F996C

The fusion of GST-HIS-Ube3C-HECT fragment was created by amplification from a plasmid template (pCAG-myc-Ube3C or respective mutants) using oligos: 5’-aagtcctctttcagggacccGTTCCATTTGAGGAGCGAG-3’ with 5’-gaattcggatcctggtacccttattaGCTCAGCTCAAAGCCAGC-3’ and inserted into the pET-49B+ expression vector (Novagen #71463) using SmaI restriction and NEBuilder system.

#### pCAG-EGFP-Ube3C

The EGFP fusion protein was created by amplification of respective templates (pRai-myc-Ube3C and pCAG-eGFP, Addgene, #89684) using the following oligos: 5’-tgtctcatcattttggcaaagATGGTGAGCAAGGGCGAGGAG-3’ with 5’-tcgaagctgaaCTTGTACAGCTCGTCCATGCC-3’ and 5’-gctgtacaagTTCAGCTTCGAAGGCGACTTC-3’ with 5’-gcggccgcgatatcctcgaggTCAGCTCAGCTCAAAGCCAGC-3’ and inserted into the pCAG-expression vector (Addgene, #11160) using EcoRI restriction and NEBuilder system.

#### pCAG-EGFP-Ube3C C1051S

The EGFP fusion protein was created by amplification of respective templates (pCAG-myc-Ube3C-C1051S and pCAG-eGFP, Addgene, #89684) using the following oligos: 5’-tgtctcatcattttggcaaagATGGTGAGCAAGGGCGAGGAG-3’ with 5’-tcgaagctgaaCTTGTACAGCTCGTCCATGCC-3’ and 5’-gctgtacaagTTCAGCTTCGAAGGCGACTTC-3’ with 5’-gcggccgcgatatcctcgaggTCAGCTCAGCTCAAAGCCAGC-3’ and inserted into the pCAG-expression vector (Addgene, #11160) using EcoRI restriction and NEBuilder system.

#### pCAG-EGFP-Ube3C S845F

The EGFP fusion protein was created by amplification of respective templates (pCAG-myc-Ube3C-S845F and pCAG-eGFP, Addgene, #89684) using the following oligos: 5’-tgtctcatcattttggcaaagATGGTGAGCAAGGGCGAGGAG-3’ with 5’-tcgaagctgaaCTTGTACAGCTCGTCCATGCC-3’ and 5’-gctgtacaagTTCAGCTTCGAAGGCGACTTC-3’ with 5’-gcggccgcgatatcctcgaggTCAGCTCAGCTCAAAGCCAGC-3’ and inserted into the pCAG-expression vector (Addgene, #11160) using EcoRI restriction and NEBuilder system.

#### pCAG-EGFP-Ube3C F996C

The EGFP fusion protein was created by amplification of respective templates (pCAG-myc-Ube3C-F996C and pCAG-eGFP, Addgene, #89684) using the following oligos: 5’-tgtctcatcattttggcaaagATGGTGAGCAAGGGCGAGGAG-3’ with 5’-tcgaagctgaaCTTGTACAGCTCGTCCATGCC-3’ and 5’-gctgtacaagTTCAGCTTCGAAGGCGACTTC-3’ with 5’-gcggccgcgatatcctcgaggTCAGCTCAGCTCAAAGCCAGC-3’inserted into the pCAG-expression vector (Addgene, #11160) using EcoRI restriction and NEBuilder system.

#### pCAG-HA-Cbll1

Mouse Cbll1 (NM_134048.2) was amplified from a P0 mouse brain cDNA library using the following oligos: 5’-gtctcatcattttggcaaagATGtacccatacgatgttccagattacgctGATCACACTGACAATG-3’ with 5’-cggccgcgatatcctcgaggTCATTGGTAATACGGTCTATATC-3’ and inserted into the pCAG-expression vector (Addgene, #11160) using EcoRI restriction and NEBuilder system.

#### pCAG-PB-eGFP

The construct was cloned via inserting the annealed oligos for the 5’ side and the 3’ site of Piggy bag into the pCAG vector. The following primers were annealed: 5’PB-ITRa: 5’-taatccctagaaagatagtctgcgtaaaattgacgcatgat-3’ with 5’PB-ITRb: 5’-taatcatgcgtcaattttacgcagactatctttctagggat-3’ and 3’PB ITRc: 5’-aattgacatgcgtcaattttacgcatgattatctttaacgtacgtcacaatatgattatctttctagggt-3’ with 3’PB-ITRd 5’-aattgccctagaaagataatcatattgtgacgtacgttaaagataatcatgcgtaaaattgacgcatgtt-3’. After annealing, the 5’ was inserted via VspI restiction side and the 3’ via MefI restriction into the pCAG-vector. Afterwards, eGFP was inserted via the EcoRI side and NEBuilder reaction using the following oligos: eGFP_fwd, 5’-gtctcatcattttggcaaagatggtgagcaagggcgag-3’ and eGFP_rev, 5’-ctagcggcgcgccaccggtgttacttgtacagctcgtccatg3’.

#### pCAG-hypPBase plasmid was a kind gift of the Jean Livet Lab

##### pCAG-PB-Ube3C

The construct was cloned by opening of pCAG-PB-flox-STOP-flox-EGFP via EcoRI and NotI. Digestion via EcoRI and NotI generated two fragments, 4360 bp and 2240 bp. The larger fragment with pCAG-PB was purified. HA-tagged Ube3C sequence was amplified using the primers: HA-msUbe3C_fwd, 5’-gtctcatcattttggcaaagATGTACCCATACGATGTTCC-3’ and HA-msUbe3C_rev, 5’-aattgacgcatgtcgcgagcTCAGCTCAGCTCAAAGCC-3’. Fragments were ligated using the NEBuilder system.

### In utero electroporation (IUE), BrdU and STM2457 pulses

IUE procedure was performed as described in our previous works and as approved by the animal welfare office ^17,26–28,71^. Briefly, DNA was diluted in TE buffer and mixed with 0.1% Fast Green FCF (Sigma-Aldrich). Final concentration of DNA used for transfecting cortical progenitors was 250 ng/μL. For the experiments with the piggyBac system, PB vectors were injected together with pCAG-hypPBase plasmid mixed 10:1 molar ratio. For the experiments described in this paper, 6 pulses of 37 V were applied using platinum electrodes. For experiments in *Ube3C* ^f/f^, we used EGFPiCre plasmid, to simultaneously express EGFP and Cre, using IRES sequence. In this plasmid, Cre sequence is modified to reduce the high CpG content of the prokaryotic coding sequence (improved Cre, iCre) to reduce a chance for epigenetic silencing. Where indicated, we used co-transfection of EGFP and iCre encoding plasmids. For the IUE experiments followed by administration of BrdU or STM2457, intraperitoneal injections were administered eight hours post-operation. For BrdU (ab142567, abcam), we used 100 μg/g in water, and for STM2457 (SML3360, Sigma-Aldrich), a single dose of 50 mg/kg in DMSO ^59^. Embryonic and early postnatal brains (until P2) were harvested and fixed in 4% PFA with 3% sucrose (v/v) overnight. Following fixation, tissue was kept in PBS until use.

### Fluorescent In Situ Hybridization (FISH)

In situ hybridization using RNAscope Technology to detect mRNA of *M. musculus Ube3C* (Probe-Mm-Ube3c-C1, 1054271-C1) was performed according to the manufacturer’s protocols (ACDBio, 323100) with a slight modification of the reduction of incubation time with the protease III to 4 min and H_2_O_2_ to 3 min. Prior to hybridization, embryonic brains at E12.5, E14.5, E16.5, and E18.5 were collected in PBS, and fixed in 4% PFA/PBS prepared with DEPC for 16–20 hours at 4 °C. Brains were then incubated in sucrose solutions (10%–20%–30%/PBS) until they reached osmotic equilibrium, embedded in O.C.T. Compound (Tissue-Tek) in a plastic cryoblock mold, and frozen on dry ice. Coronal sections with a thickness of 16 μm were collected using Leica CM3050S or RWD Minus FS800 cryostat.

### Antibodies

Primary antibodies used for Western blotting in this publication were used at a dilution 1:750 in 3% BSA in TBS-T buffer with 0.1% Tween-20, apart from anti-β-actin, used at 1:2 000. Secondary antibodies coupled with HRP were diluted 1:10 000 in the same buffer as the primary ones. For immunohistochemistry, primary antibodies and the respective fluorophore-linked secondary ones were diluted 1:500 in the blocking buffer. For ubiquitination assays or co-immunoprecipitation, the antibodies and working conditions are indicated in the respective sections below. Detailed list of critical reagents contains specification of primary and secondary antibodies used in this study.

### Cryosectioning

For histological procedures, brain sections were prepared on Leica CM3050S and RWD Minux FS800 cryostat. Fixed brains were incubated with sucrose gradient prior to freezing. Next, brains were flash-frozen in −38 to −40 °C isopentane (Roth), or frozen on dry ice as cryoblocks, after embedding in Tissue-Tek OCT compound (Sakura). Free-floating coronal cryosections of 50 μm thickness were collected in PBS/0.01% sodium azide solution. For in situ hybridization, BrdU birthdating, or the experiments with STM2457, 12-16 μm thickness sections were directly collected on the Superfrost glass slides.

### Immunohistochemistry

Fixed brain sections were washed with PBS three times at room temperature prior to the procedure to remove the sucrose and freezing compound residue. The sections were then incubated with the blocking solution for one hour at room temperature, next with primary antibody and diluted DAPI for 16-20 hours at 4°C. Sections were then washed in PBS three times for up to 30 min and incubated with secondary antibody diluted in the blocking buffer for up to four hours at room temperature, 1:250. After three washes with PBS for up to 30 minutes, sections were mounted with cover glass (Menzel-Gläser) and Immu-Mount mounting medium (Shandon, Thermo-Scientific). For the experiments with BrdU ^17,72^ and m6A, cryosections were let dry overnight at 4°C, briefly washed in PBS and incubated with boiling citrate-based Antigen Unmasking Solution (Vector, pH 6.0) for ten minutes in a microwave. After cooling on ice, sections were further processed for immunohistochemistry. Blocking solution: 5% horse serum (Corning), 0.5% (v/v) Triton X-100 (Sigma-Aldrich), PBS.

### Biochemical experiments

Details of the protocols used for sodium dodecyl sulfate polyacrylamide gel electrophoresis (SDS-PAGE) and Western blotting were performed according to standard lab practice and are thoroughly described in our previous works ^17,27,28,73^. Nitrocellulose membranes were used for Western blotting. Acquisition of the chemiluminescence for the biochemical experiments in this manuscript was performed using BioRad Molecular Imager, ChemiDoc XRS+ and Azure Biosystems 600.

### Preparation of cortical lysates

Unless explicitly stated otherwise, for quantitative biochemistry, cortical tissue was dissected on ice and immediately snap-frozen in liquid nitrogen until further processing. Tissue was lysed and homogenized in Buffer A, supplemented with 2.5 mM MgCl2, 0.025 U/µl benzonase (70664-3, Merck), complete protease inhibitor cocktail (11873580001, Roche), MG-132 (Sigma-Aldrich), and NEM (E3876, Sigma-Aldrich). After homogenization, cortices were incubated for 20 min at 70°C with the following centrifugation for 15 min at 12,000 g. Protein concentration in the supernatant was determined using the BCA method (Thermo, Pierce), or DC protein assay (Bio-Rad).

Buffer A: 50 mM Tris-Cl pH 7.5, 300 mM NaCl, 1% SDS, 0.2 mM PMSF, 10 mM NEM, 20 µM MG-132, Benzonase, 20 mM MgCl_2_, complete protease inhibitor

### Structure modeling

For modeling of Ube3C critical residues within its structure, we used AlphaFold 3 ^74^.

### In vivo ubiquitination assay (endogenous immunoprecipitation in denaturing conditions)

Frozen P0 cortices were lysed and homogenized (mechanical dissociation by pipetting) in 400 µL Buffer A. Lysed samples were then incubated at 95°C for 5 min. Remaining supernatant was diluted 1:10 with Buffer B and centrifuged at 12,000 g at 4°C for 20 minutes. Protein G Sepharose beads (17-0618-01, GE Healthcare) were washed three times in Buffer B. The supernatant of each sample was added to washed, unconjugated beads and incubated for a minimum of 1 hour at room temperature, rotating. Primary antibody (rabbit anti-Cbll1, A302-969A-M, Bethyl) or normal rabbit IgG (#2729, CST) was added to 50 µL bead slurry/assay point (per one P0 cortex) and coupled for a minimum of 1 hour at room temperature, rotating. Pre-cleared supernatants and coupled beads were incubated for one hour at room temperature, rotating. The beads were then washed three times, incubated with 50 µL Lämmli buffer at 70°C for 20 min and further analyzed for ubiquitin mark using Western blotting. To visualize ubiquitination, we used rabbit anti-ubiquitin (E4I2J) antibody (43124, CST).

Buffer A: 50 mM Tris-Cl pH 7.5, 300 mM NaCl, 1% SDS, 0.2 mM PMSF, 10 mM NEM, 20 µM MG-132, Benzonase, 20 mM MgCl_2_, complete protease inhibitor Buffer B: 50 mM Tris-Cl pH 7.5, 300 mM NaCl, 1% Tritox-100, 0.2 mM PMSF, 10 mM NEM, 20 µM MG-132, Benzonase, 20 mM MgCl_2_, complete protease inhibitor

### Protein purification

Ube3C HECT fragments and its variants were expressed in *E. coli* LOBSTR-BL21(DE3)-RIL (Kerafast) grown in Terrific Broth (TB), induced with 0.5 mM IPTG at 15 °C overnight. Cells were lysed in 50 mM Tris pH 7.5, 200 mM NaCl, 5 mM DTT, 4 mM MgCl2 and the soluble lysate fraction incubated with Glutathione Sepharose 4B resin (Cytiva) for 2 hours at 4 °C. The resin was washed 3 times with 10 column volumes of lysis buffer and resuspended as a 50% slurry. Ube3C was liberated from the GST-tag by protease digestion with HRV-3C (prepared in-house) at 4 °C overnight before subjected to size-exclusion chromatography (SEC) in 10 mM Tris pH 7.8, 150 mM NaCl, 4 mM MgCl_2_, 2 mM TCEP using a HiLoad Superdex 200 16/600column (Cytiva).

### In vitro autoubiquitination assay

Autoubiquitination activity of recombinant WT or variant Ube3C HECT was measured using an in vitro reconstitution system. Recombinant proteins were expressed using a pET vector system in E. coli, using standard protocols, according to our previous works ^75,76^. While assembling the reaction, all components were kept on ice. Reactions of 205.6 μl total volume were prepared by adding reagents in the following order to contain; 15 μM Ub (U6253, Merck), 1 μM E2 enzyme (UBCH5C), and 3 μM or 6 μM of the corresponding Ube3C HECT variant. Reactions were filled to 200 μl with reaction buffer (10 mM Tris-HCl, pH 7.8, 50 mM NaCl, 4 mM MgCl_2_, 8 mM ATP). The reaction was initiated by adding 0.05 μM E1 enzyme (UBA1) and incubated at 37°C. 30 μL samples were withdrawn at 1, 5, 10, 30 and 60 min, and quenched in 10 μL Lämmli sample buffer. Samples were denatured at 95°C for 5 min and resolved by SDS-PAGE. 10-20 μL sample and 5 μL PageRuler™ Plus Prestained Protein Ladder (Thermo-Fisher) were loaded into 4-20% Tris-Glycine gradient gels (XP0420C, Thermo-Fisher) and placed in an Invitrogen mini gel tank filled with Tris-Glycine SDS running buffer. Gels were resolved at 100 V for approximately one hour before staining with Quick Coomassie Stain (NB-45-00078, Neo-Biotech) overnight. Gels were transferred to water and scanned with an Epson Perfection V39 scanner. Densitometry was performed using Fiji software to quantify the intensity corresponding to free ubiquitin at 5 min reaction timepoint. Each time point was expressed as a percentage of the signal at 0 min.

### Differential Scanning Fluorimetry (DSF)

DSF assay was carried out using Ube3C HECTs, with their validated activity using an autoubiquitination assay (Fig. 2b) comprising variants at the final concentration of 1 µM in the assay buffer composed of 1 µM UbcH7, 0.1 µM E1 enzyme, 5 mM ATP, 80 µM ubiquitin, and 20 µM ubiquitin CW800. DSF measurements were performed with a CFX96 Real-Time System (C1000 Thermal Cycler, Bio-Rad) in combination with SYPRO-Orange protein stain (Invitrogen). Purified Ube3C at 9 µM was mixed with 5X SYPRO-Orange in protein purification buffer (10 mM Tris pH 7.8, 150 mM NaCl, 4 mM MgCl_2_, 2 mM TCEP). Samples were incubated at 20 °C for 5 minutes, before increasing to 95 °C in 0.5 -°C increments with 45 sec incubations. Protein unfolding at each temperature step was monitored by SYPRO-orange fluorescence in the HEX channel. Data were collected in triplicate, processed using MoltenProt (https://spc.embl-hamburg.de/app/moltenprot; ^77,78^).

### Cell-based in vitro ubiquitination assay

Ubiquitination of Cbll1 by Ube3C was assayed in HEK293T cells transfected using Lipofectamine 2000 (11668019, Thermo-Fisher) according to published protocols ^79,80^. Cells were collected in PBS after 16-20 hours of expression in culture using centrifugation at 800 g for 5 min at 4°C. After sampling the input, cells were lysed in 1 mL of buffer A, supplemented with benzonase (1 µL/1 mL; 70746, Merck) and 10 mM MgCl_2_ and subjected to sonification using three cycles with 5 strokes and 10 sec breaks. The supernatant was then collected after centrifugation for 10 min at 12 000 g at 4°C and incubated with 50 μL equilibrated Ni-NTA-agarose bead slurry (Ni Sepharose 6 Fast Flow, 17531801, Cytiva) for 16-20 hours at 4°C. Beads were washed sequentially with 1 mL of buffer A, buffer A and B 1:1 and buffer B, dried, incubated with 50 μL of sample Lämmli buffer and imidazole (2X / 200 mM) at 95°C for 10 min, and further processed for Western blotting. For the analysis, we used the following antibodies at indicated dilutions in 3% BSA/1X TBS/0.1% Tween-20 blocking buffer: mouse anti-His (HIS.H8, ab18184, abcam), 1:1000; mouse anti-myc (9B11, 2276, CST), 1:300; rabbit anti-myc (2278, CST), 1:300; and rabbit anti-HA (3724, CST), 1:1000.

Buffer A: 6 M guanidin-HCl, 100 mM phosphate buffer (pH 8.0), 10 mM imidazol, 0.4% Triton X-100, 0.125 M NaCl, supplemented with NEM (Sigma-Aldrich), MG132 (Sigma-Aldrich), and protease inhibitors (Roche) Buffer B: 25 mM Tris-HCl (pH 6.8), 20 mM imidazol

### Co-Immunoprecipitation (Co-IP)

Experiments for the interaction between the EGFP-Ube3C fusion variants and HA-Cbll1 were performed in HEK293T cells, seeded in 6 cm dishes, after transfection using Lipofectamine 2000 (11668019, Thermo-Fisher). After 16-20 hours of expression, cells were collected by scraping the bottom of each dish cells and centrifugation at 800 g at 4°C. Cells from one well (one assay point) were lysed in 1 mL Co-IP lysis buffer, subjected to sonification (four times, 5 sec, with 10 sec break at 20% power) and centrifuged 10 min at 12 000 g at 4°C. Supernatant was incubated with equilibrated 50 µL GFP bead slurry (ChromoTek GFP-Trap Agarose, GTA) per assay point for two hours at 4°C. Beads were then washed once with the Co-IP lysis buffer, three times with wash buffer, dried and incubated with 60 µL Lämmli sample buffer in TBS at 95°C for 10 min. For Western blotting, we used the following antibodies with indicated dilutions in 3% BSA / 1X TBS / 0.1% Tween-20 blocking buffer: goat anti-EGFP (600-101-215, Rockland), 1:1000; rabbit anti-HA (3724, CST), 1:1000.

Co-IP lysis buffer: 10 mM Tris-Cl (pH 8.0), 150 mM NaCl, 10 mM MgCl_2_, 1% Triton X-100, 0.1% SDS, 0.1% sodium deoxycholate, benzonase (1 µL/1 mL; 70746, Merck), protease cocktail inhibitor (Roche) Wash buffer: 10 mM Tris-Cl (pH 8.0), 150 mM NaCl, 0.5% Triton X-100 TBS (1X): 50 mM Tris-Cl (pH 7.5), 150 mM NaCl

### TUBE proteomics sample preparation

To identify ubiquitomes in control and *Ube3C* cKO E14.5 cortices, we used TUBE technology (Tebu-bio; UM407M and UM0500M). We prepared three independent biological replicates per genotype. One cortex was isolated in PBS on ice under the dissecting microscope and snap frozen in liquid nitrogen until batch processing. Per cortex, we used 200 µL TUBE lysis buffer and homogenization using 1 mL syringe, with 30 µL of supernatant reserved as input. Homogenates were clarified using centrifugation at 12 000 g for 20 min at 4°C. An 80 µL of magnetic bead slurry was equilibrated in TBS-T buffer and incubated with the clarified lysate for 2 hours at 4°C. Beads were then washed with wash buffer 1 and 2 and either eluted by incubation with 40 µL Lämmli sample buffer for 20 min at 70°C for the analysis with Western blotting, or washed once more with 100 mM Tris-HCl, pH7.5 and analyzed further by affinity proteomics mass spectrometry.

TUBE lysis buffer: 100 mM Tris-HCl (pH 8.0), 150 mM NaCl, 5 mM EDTA, 1% NP-40, PR619 (SML0430, Merck), 5 mM NEM (Merck).

Wash buffer 1: 20 mM Tris-HCl (pH 7.5), 250 mM NaCl, 0.2% NP-40, 1 mM DTT

Wash buffer 2: 20 mM Tris-HCl (pH 7.5), 150 mM NaCl, 0.05% NP-40, 1 mM DTT

TBS-T buffer: 20 mM Tris-HCl (pH 8.0), 150 mM NaCl, 0.1% Tween-20

### Control- and K48-TUBE affinity proteomics mass spectrometry

Pulldown beads were resuspended in digestion buffer (6 M urea / 2 M thiourea in 50 mM ammonium bicarbonate), reduced with dithiothreitol (10 mM final concentration) for 30 minutes, followed by alkylation with chloroacetamide (55 mM final concentration) for 45 minutes. Samples were diluted with four volumes 50 mM ammonium bicarbonate and incubated over night at room temperature with 500 ng Endopeptidase LysC (Wako, Neuss) and 1 µL sequence grade trypsin (Promega, Mannheim, GER). After adding formic acid (final concentration 1%), peptides were desalted and separated by reversed phase chromatography on an in-house manufactured 20 cm fritless silica microcolumns (inner diameter of 75 µm, packed with ReproSil-Pur C18-AQ 1.9 µm resin (Dr. Maisch GmbH)) using a 98-min gradient of increasing Buffer B (90% ACN, 0.1% FA) concentration (from 2% to 60%) with a 250 nl/min flow rate on a High-Performance Liquid Chromatography (HPLC) system (Thermo Fisher Scientific). Eluting peptides were directly ionized by electrospray ionization and transferred into a HF-X mass spectrometer (Thermo Fisher Scientific), which was operated in the data-dependent mode with performing full scans (60K resolution; 3 x 106 ion count target; maximum injection time 10 ms), followed by top 20 MS2 scans at 15K resolution with a maximum injection time of 100ms. Only precursor with charge states between 2-7 were fragmented. Dynamic exclusion was set to 30 sec. Raw data were analyzed using the MaxQuant software (v1.6.3.4) with UniProt mouse database (MOUSE.2022-03). The search included variable modifications of methionine oxidation and N-terminal acetylation, deamidation (N and Q) and fixed modification of carbamidomethyl cysteine. The FDR was set to 1% for peptide and protein identifications. The integrated label-free quantification algorithm was activated. Retention times were recalibrated based on the built-in nonlinear time-rescaling algorithm and MS/MS identifications were transferred between LC-MS/MS runs with the “Match between runs” option. LFQ intensity values were used for quantification. Reverse hits, contaminants and proteins only identified by site were filtered out. Replicates for each condition were defined as groups and intensity values were filtered for “minimum value of 3” per group. After log2 transformation, missing values were imputed with random noise simulating the detection limit of the mass spectrometer. Differential protein abundance was calculated using two-sample Student’s t test. Differential proteins were defined based on FDR 5%. These experiments were performed by MK and PM.

### Global Proteome Profiling with TMT

For proteomic profiling by mass spectrometry, tissue samples were cryopulverized, suspended in lysis buffer (1% Sodium deoxcholate, 100 mM Tris-HCl pH 8, 150 mM NaCl, 1 mM EDTA, 40 mM CAA, 10mM DTT), heated for 10 minutes at 95°C, cooled to room temperature and treated for 15 minutes with benzonase (Merck, 50 units) for 30 min at 37°C. Endopeptidase LysC (Wako) and sequence-grade trypsin (Promega) were added to the samples with an enzyme-to-protein ratio of 1:50, followed by an incubation over night at 37°C. Peptides were desalted, resuspended in 50 mM HEPES and labeled with 16-plex tandem mass tag reagents (TMTpro, Fisher Scientific) following the vendors instructions. Samples for each plex were combined, desalted and fractionated by high-pH reversed phase off-line chromatography (1290 Infinity, Agilent) and pooled into 30 fractions. For LC-MS/MS measurements, peptides were reconstituted in 3% acetonitrile with 0.1% formic acid and separated on a reversed-phase column (20 cm fritless silica microcolumns (inner diameter of 75 µm, packed with ReproSil-Pur C18-AQ 1.9 µm resin (Dr. Maisch GmbH)), using a 98-min gradient of increasing Buffer B (90% ACN, 0.1% FA) concentration (from 2% to 60%) with a 250 nl/min flow rate on a High-Performance Liquid Chromatography (HPLC) system (Thermo Fisher Scientific) and analyzed on an Exploris 480 instrument (Thermo Fisher Scientific). The mass spectrometer was operated in data-dependent acquisition mode using 60K resolution, 350–1500 m/z scan range, maximum injection time of 50 ms. The top 20 MS/MS scan were obtained at 45K resolution with 0.4 m/z isolation window and a maximum injection time of 86 ms. Dynamic exclusion was set to 20 s and only precursor with a charge state between 2-6 were selected for fragmentation. RAW data were analyzed with MaxQuant software package (v 1.6.10.43) using the Uniprot mouse databases (MOUSE_2019_07). The search included variable modifications of methionine oxidation and N-terminal acetylation, deamidation (N and Q) and fixed modification of carbamidomethyl cysteine. Reporter ion MS2 for TMT16 was selected (internal and N-terminal) and TMT batch specific corrections factors were specified. The FDR (false discovery rate) was set to 1% for peptide and protein identifications. The resulting text files were filtered to exclude reverse database hits and potential contaminants. Only proteins detected in all samples of each plex were used for quantitation. Sample intensities were log2 transformed, normalized to the reference channel (mixture of all samples) and z-scored within each sample. Differences in protein abundance between experimental groups were calculated using Student’s T-Test. Signals passing the significance cut-off of FDR 10% were considered differentially expressed. These experiments were performed by MK and PM.

### Endogenous Cbll1 immunoprecipitation

For the detection of embryonic cortex-specific Cbll1 interactome, we used IgG- and anti-Cbll1-coupled (A302-969A-M, Bethyl) G-Protein Sepharose (P3296, Merck) beads. For mass spectrometry analysis, we subjected three biological replicates per condition, each made up of two pooled dissected neocortices. Per assay point, we used 2 mL of lysis buffer. Each sample was centrifugated for 20 min at 12 000 g and 4°C and the supernatant pre-cleared by incubation with 25 µL equilibrated Sepharose bead slurry for 1 hour at 4°C. For the immunoprecipitation, we used 50 µL bead slurry per assay point, coupled with 1 µL of either IgG or anti-Cbll1 antibody (each 1 mg/mL), for 1 hour at 4°C. The immunoglobulin-coupled beads were incubated with the pre-cleared supernatant at 4°C for 16-20 hours, then beads were washed with the wash buffer and incubated either with Lämmli sample buffer for Western blot validation or proteins bound to the beads were eluted in 100 mM HEPES (pH 8.0), 1% SDS and subjected to mass spectrometry.

Lysis buffer: 50 mM Tris-HCl (pH 7.5), 300 mM NaCl, 1% Triton X-100, supplemented with benzonase (1 µL/1 mL; 70746, Merck), 10 mM MgCl2, and protease inhibitors (Roche) Wash buffer: 25 mM Tris-HCl (pH 7.5), 150 mM NaCl, 0.5% Triton X-100

### Proteomic identification of Cbll1 interactome in vivo

For the mass spectrometric analysis of Cbll1/Hakai enriched samples in-solution tryptic digest were performed following a modified version of the Single-Pot Solid-Phase-enhanced Sample Preparation (SP3) technology ^81,82^. 20 µL of a slurry of hydrophilic and hydrophobic Sera-Mag Beads (Thermo Scientific, #4515-2105-050250, 6515-2105-050250) were mixed, washed with water and were then reconstituted in 100 µL water. 5 µl of the prepared bead slurry were added to 50 µL of the eluate following the addition of 55 µL of acetonitrile. All further steps were prepared using the King Fisher Apex System (Thermo Scientific). After binding to beads, beads were washed three times with 100 µl of 80% ethanol before they were transferred to 100 µL of digestion buffer (50 mM Hepes/NaOH pH 8.4 supplemented with 5 mM TCEP, 20 mM chloroacetamide (Sigma-Aldrich, #C0267), and 0.25 µg trypsin (Promega, #V5111)). Samples were digested over night at 37°C, beads were removed and the remaining peptides were dried down and subsequently reconstituted in 10 µL of water. 80 µg of TMT6plex (Thermo Scientific, #90066) ^83^ label reagent dissolved in 4 µL of acetonitrile were added and the mixture was incubated for 1 hour at room temperature (IgG control: TMT126, TMT127N, TMT128C (n=3); Hakai: TMT129N, TMT130C, TMT131N (n=3)). Excess TMT reagent was quenched by the addition of 4 µL of an aqueous solution of 5% hydroxylamine (Sigma, 438227). Mixed peptides were subjected to a reverse phase clean-up step (OASIS HLB 96-well µElution Plate, Waters #186001828BA). Peptides were analyzed on analyzed by LC-MS/MS on an Orbitrap Fusion Lumos mass spectrometer (Thermo Scentific). To this end, peptides were separated using an Ultimate 3000 nano RSLC system (Dionex) equipped with a trapping cartridge (Precolumn C18 PepMap100, 5 mm, 300 μm i.d., 5 μm, 100 Å) and an analytical column (Acclaim PepMap 100. 75 × 50 cm C18, 3 mm, 100 Å) connected to a nanospray-Flex ion source. The peptides were loaded onto the trap column at 30 µL per min using solvent A (0.1% formic acid) and eluted using a gradient from 2 to 38% Solvent B (0.1% formic acid in acetonitrile) over 90 min at 0.3 µL per min (all solvents were of LC-MS grade). The Orbitrap Fusion Lumos was operated in positive ion mode with a spray voltage of 2.4 kV and capillary temperature of 275 °C. Full scan MS spectra with a mass range of 375–1500 m/z were acquired in profile mode using a resolution of 60,000 (maximum fill time of 50 ms; AGC Target was set to Standard) and a RF lens setting of 30%. Fragmentation was triggered for 3 s cycle time for peptide like features with charge states of 2–7 on the MS scan (data-dependent acquisition). Precursors were isolated using the quadrupole with a window of 0.7 m/z and fragmented with a normalized collision energy of 36%. Fragment mass spectra were acquired in profile mode and an orbitrap resolution of 15,000. Maximum fill time was set to 54 ms. AGC target was set to 200%. The dynamic exclusion was set to 60 s. Acquired data were analyzed using FragPipe (PMID: 28394336) and a Uniprot *Mus musculus* fasts database (UP000000589, ID10090 with 21.968 entries, date: 27.10.2022, downloaded: January 11^th^ 2023) including common contaminants. The following modifications were considered: Carbamidomethyl (C, fixed), TMT6plex (K, fixed), Acetyl (N-term, variable), Oxidation (M, variable) and TMT6plex (N-term, variable). The mass error tolerance for full scan MS spectra was set to 10 ppm and for MS/MS spectra to 0.02 Da. A maximum of 2 missed cleavages were allowed. A minimum of 2 unique peptides with a peptide length of at least seven amino acids and a false discovery rate below 0.01 were required on the peptide and protein level ^84^. The raw output files of FragPipe (protein.tsv – files ^85^) were processed using the R programming language (ISBN 3-900051-07-0). Contaminants were filtered out and only proteins that were quantified with at least two razor peptides were considered for the analysis. 1072 proteins passed the quality control filters. Log2 transformed raw TMT reporter ion intensities were first cleaned for batch effects using the ‘removeBatchEffects’ function of the limma package ^86^ and further normalized using the vsn package (variance stabilization normalization ^87^). Proteins were tested for differential expression using the limma package. The replicate information was added as a factor in the design matrix given as an argument to the ‘lmFit’ function of limma. A protein was annotated as a hit with a false discovery rate (fdr) smaller 5 % and a fold-change of at least 100 % and as a candidate with a fdr below 20 % and a fold-change of at least 50 %. This experiment was performed by FS and PH.

### Gene Set Enrichment Analysis (GSEA)

GSEA was performed using an open source package https://bioinformatics.sdstate.edu/go/ using GO ontology database https://zenodo.org/records/14861039 ^88^. For proteomics data (Fig. 4c and 6b), our input represented proteins with lower ubiquitination or higher protein levels in *Ube3C* cKO, respectively; for Fig. 5g-5i, we used all significantly changed transcripts.

### iPSC line culture

The human iPS cell line TMOi001-A (A18945, Thermo Fisher Scientific) was cultured in standard hypoxic conditions (37°C, 5% CO2, 5% O2) in E8 Flex medium (Thermo Fisher Scientific) on Geltrex-coated plates (Thermo Fisher Scientifc).

### Generation of *UBE3C* Knockout iPSCs

*UBE3C* knock-out iPSCs were generated using the Alt-R CRISPR/Cas9 system (IDT). In short, sgRNAs targeting the CDS of the UBE3C gene were designed and ranked with the CRISpick tool ^89,90^ and a single sgRNA targeting exon 7 of the *UBE3C* gene (5’-ACTTACAACTCCTGTCCGGA-3’) selected for its minimal off-target activity. For the transfection, iPSCs were dissociated into single cells using TrypLE (Thermo Fisher Scientific) and nucleofected with pre-assembled sgRNA-Cas9 RNP complex using the P3 Primary Cell 4D-Nucleofector X Kit (Lonza, V4XP-3032). On day 4 post-transfection, iPSCs were dissociated into a single-cell suspension and seeded onto a miniaturized low-volume chamber grid for automated single-cell cloning (IsoCell platform, iota Sciences) as described previously ^91^. Half an hour post-plating, monoclonal wells were identified and expanded for 10 days. Expanded single-cell clones were transferred onto 2 96-well plates (1:1) and cultured until they reached 70-80% confluency. Single-cell clones with desired homozygous out-of-frame indel modifications were identified with Sanger sequencing and further expanded for validation and biobanking.

### Organoids generation

The organoids were generated using forebrain protocols following a modified protocol ^92^. Briefly, after dissociation into a single-cell suspension using TrypLE, 9,000 cells were seeded per well in 96-well plates with 100 µL of E8 medium containing 50 µM ROCK inhibitor. The next day, 80% of the medium was replaced with E6 medium supplemented with 0.1 µM LDN, 5 µM XAV, and 10 µM SB. On day 5, the medium was replaced with E6 medium supplemented only with LDN and SB (dual SMAD inhibition). Between days 7 and 9, liquid embedding of the organoids was performed using 2% Matrigel (Corning, 356234) in organoid differentiation medium. This medium consisted of a 1:1 mixture of DMEM/F12 and Neurobasal, supplemented with 1× N2 supplement, 1× B27 - vitamin A supplement, insulin, 2-ME solution, GlutaMAX, MEM-NEAA, and LIF. The embedded organoids were cultured in ultra-low-attachment 6-well plates on an orbital shaker (80 rpm) until day 15. Subsequently, the medium was replaced with organoid maturation medium, consisting of a 1:1 mixture of DMEM/F12 and Neurobasal, supplemented with N2 supplement, B27 + vitamin A supplement, insulin, 2-ME solution, GlutaMAX, MEM-NEAA, sodium bicarbonate, vitamin C solution, chemically defined lipid concentrate, BDNF, GDNF, and cAMP. Organoids were collected at day 5 (5D), 15 and 30 for analyses.

### Immunostaining of brain organoids and image acquisition

For immunostainings, organoids were washed three times with PBS and fixed in 4% paraformaldehyde for 20 to 60 min (depending on the organoid size) at 4°C, then washed with PBS three times for 10 min each. The tissue was incubated in 40% sucrose (in PBS) until it sunk (overnight) and then embedded in 13%/10% gelatin/sucrose. Frozen blocks were stored at -80 °C, prior to cryosections. 10-12 μm sections were prepared using a cryostat. Sections were incubated with warm PBS for 10-15 min in order to remove the embedding medium and then fixed for additional 10 min with 4% PFA, washed three times with PBS and blocked and permeabilized in 0.25% Triton-X, 5% normal goat serum in PBS for 1 h. Sections were first incubated with primary antibodies in 0.1% Triton-X, 5% normal goat serum overnight. They were then washed three times for 10 min each with PBS-T (0.1% Triton X-100) and incubated with secondary antibodies at RT for 2 h, followed by staining with DAPI (final 1 µg/ml) for 10 min and washed three times with PBS-T. The images were acquired using a Keyence BZ-X710 (Osaka, Japan) microscope or confocal microscopes specified in the Image acquisition section of the Methods. Organoids were generated and analysed by ARW, IK, and NK.

### Single-nuclei RNA sequencing (sn-RNAseq)

For nuclei isolation, frozen organoids were disrupted with 10-15 strokes of a plastic pestle in 500 µL of NP-40 lysis buffer (10 mM Tris-HCl (pH 7.4), 10 mM NaCl, 5.6 mM MgCl_2_, 0.1% NP-40, 1 mM DTT, Complete EDTA-free protease inhibitor, 2% BSA, 1 U/µl Protector RNase inhibitor) and incubated on ice for 5 minutes, with gentle pipette mixing after 2.5 minutes. The nuclei suspension was then filtered through a 70 µm strainer into a fresh 1,5 mL Eppendorf tube and nuclei were sedimented by centrifugation at 500 g for 5 min at 4°C. After removing the supernatant, 1 mL of cold wash buffer (PBS, 1% BSA, 0.4 U/µl Takara RNAse inhibitor) was added without disturbing the pellet. Nuclei were incubated on ice for 5 minutes and then gently resuspended and sedimented again at 500g for 5 min at 4°C. Nuclei were then resuspended in 300-500 µL of cold wash buffer and filtered through a 40 µm Flowmi cell strainer. After adding DAPI (final concentration 0,1 µg/mL) nuclei were sorted on an Aria III instrument with a 100 µm nozzle in an Eppendorf tube containing 100 µL sort buffer (PBS, 2% BSA, 2.5 U/µl Protector RNAse inhibitor). Single nucleus RNA sequencing libraries were prepared using the Chromium Single Cell 3’ GEM-X v4 assay (10X Genomics). In short, cell suspensions were encapsulated into Gel Beads in Emulsion (GEMs) using the Chromium Controller. Within each GEM, cell lysis and barcoded reverse transcription of RNA occurred, followed by cDNA amplification. The amplified cDNA underwent library construction via fragmentation, end-repair, A-tailing, adaptor ligation, and index PCR. Final libraries were sequenced on an Illumina NovaSeq 6000 system to a minimum depth of 20.000 reads per cell. These experiments were performed by CQ, TB, CD and TC.

### scRNA-seq data analysis

We processed demultiplex, raw paired-end snRNAseq data using Cell Ranger (v. 8.1.0) from 10x Genomics (https://www.10xgenomics.com/support/software/cell-ranger/latest). We used the “cell ranger count” function to align reads to the human genome (GRCh38) and generate digital gene expression (DGE) matrices for each sample. DGEs were further processed in R (v. 4.4.1) using Seurat ^93^ (v. 5.1.0). Raw DGEs were imported and only genes detected in at least 5 cells were kept for downstream analyses. Similarly, cells will less than 2000 unique transcripts were discarded. After merging all samples in a single Seurat object, we performed gene expression normalization and scaling using SCTransform ^94^. We then performed dimensionality reduction and selected the first 10 principal components for shared nearest neighbours identification, clustering (resolution 0.5) and computing a two-dimensional UMAP embeddings. We removed two control clusters (clusters 3 and 14) and four clusters across all organoids (clusters 0 and 10) that were characterized by a substantially lower UMI counts. Clusters were manually annotated in 3 main cell types analyzing the expression of canonical marker genes. To identify differentially regulated genes upon *UBE3C* knock-out in robust and reproducible way, we performed a pseudobulk analysis using the DESeq2 package ^95^ (v. 1.44.0). For each celltype, we generated a pseudobulk expression profile for each organoid by summing the expression of 1500 randomly selected cells (with replacement). We then filtered genes with less 50 counts in less than 2 samples expression values and comparesd gene expression between KO and CTR groups. For each comparison, we identified differentially expressed genes and adjusted *P* value was calculated using Bonferroni correction for multiple testing correction. The analysis was performed by TP.

### Quantification of m6A in control and *Ube3C* cKO samples

We assayed the relative amounts of m6A in the *Ube3C* cKO neocortices using three different methods. We performed a colorimetric assay using a commercially available kit, EpiQuik m6A RNA Methylation Quantification Kit (P-9005, EpigenTek). Prior to the analysis, RNA was isolated using the Trizol (Thermo-Fisher) extraction method. E14.5 and P0 cortical tissue was shredded using Ultra-Turrax in 1 mL Trizol. After extraction with chloroform, RNA was pelleted using ethanol, washed, let dry and resuspended in ddH_2_O. Quality of the RNA prep was determined via A260/280 ratio. For the samples used in our experiments, the ratio was between 2.00 and 2.15. We quantified the total m6A and %m6A in the RNA using SpectraMax iD3 (Molecular Devices) plate reader. Next, we performed a dot blot assay to detect m6A in isolated RNA. We followed the manufacturer’s protocol (m6A Dot Blot Assay Kit, DB-m6A, RayBiotech). Per dot, we applied 300 ng RNA in 1 µL. Density of the m6A signal was normalized to the total amount of RNA as visualized by Methylene Blue stain and expressed as fraction of control per developmental stage. For the m6A immunohistochemistry, we used thin 16 µm cryosections and antigen retrieval, as described for the BrdU analysis. Images were then binarized and particle size and numbers were quantified using Analyze Particles command in Fiji, after Watershed Segmentaion. For each experiment, a matched number of cortices of each sex were analyzed and the measurements pooled.

### Image acquisition

Brain sections after immunohistochemical processing were imaged using Zeiss Spinning Disc microscope, TCS SL upright, or Zeiss LSM900 Airyscan2 confocal.

### Quantitative Western blotting and dot blot analysis

Quantification of protein levels was performed with ImageStudioLite (Li-Cor) software after exposure of nitrocellulose membranes to ECL chemiluminescence with signals acquired by Bio-Rad Molecular Imager, ChemiDoc XRS+, or Azure Biosystems 600. For the image analysis, we used ImageLab, ImageStudioLite and Fiji software. Protein level was expressed relative to the signal for β-actin or vinculin detected in respective lane or to the total protein as measured by the integral of Coomassie Staining intensity (NB-45-00078, Neo-Biotech), measured with Tracing tool of ImageJ software.

### Cortical lamination and colocalization with marker proteins

We based our positioning analysis on our previous works ^17,27^. To determine the position of neurons in the cortical plate, confocal images of EGFP signals were first transformed so that the pia is perpendicularly oriented to the horizontal axis. Next, the positions of neuronal somata were marked in Fiji (Cell Counter plugin). Using the y-coordinate, we then expressed the position of a given cell relative to the size of the CP and plotted it as % CP, with 0% representing the bottom of the cortical plate / the subplate (SP) and a 100% the pial surface / the marginal zone (MZ). Positions of all cells were then plotted individually on the graph. Only brains with comparable electroporation efficiencies were analyzed. We also provided the normalized quantification using cortical bins. For this, the number of neurons in each of the five cortical bins was normalized by the total number of electroporated neurons within a given cortical section. For colocalization with marker proteins, we manually determined the expression of analyzed protein based on the immunohistochemical signals in cryosections.

### Statistical analysis and reproducibility

All statistics were computed using Prism Graph Pad software (version 10.1.0, 264, released Oct 18, 2023 and previous versions). Description of statistical tests, definition of center, dispersion, precision, and definition of significance are listed in the figure legends. Briefly, the distribution of data points was determined using normality tests (D’Agostino-Pearson, Anderson-Darling, Shapiro-Wilk, or Kolmogorov-Smirnov). We used unpaired two-tailed t-test, or one/two-way ANOVA with respective post hoc test; Mann-Whitney test to compare between two groups, or Kruskal-Wallis test with Dunn’s multiple comparison test for comparisons between multiple groups. Western blotting, biochemical assays, as well as IUE, immunohistochemistry or FISH presented in this manuscript were repeated at least three times. Representative micrographs or blots are derived from the analyzed datasets.

## CRITICAL REGAENTS

CRITICAL REAGENTS
| REAGENT or RESOURCE | SOURCE | IDENTIFIER |
| --- | --- | --- |
| **Primary Antibodies** |  |  |
| Rabbit polyclonal anti-Sox9 | Millipore | AB5535; RRID: AB_2239761 |
| Mouse monoclonal anti-Satb2 | SCBT | sc-81376; RRID: AB_1129287 |
| Rabbit polyclonal anti-Satb2 | Self-made <sup>96</sup> | N/A |
| Mouse monoclonal anti-β actin | Millipore | MABT825; RRID: AB_2571580 |
| Rabbit polyclonal anti-Cbll1 | Bethyl | A302-969A; RRID: AB_10754975 |
| Rabbit polyclonal anti-Ube3C | Abcam | ab243830 |
| Rabbit polyclonal anti-Ube3C N1N3 | GeneTex | GTX119102; RRID: AB_10617623 |
| Goat polyclonal anti-EGFP | Rockland | 600-101-215; RRID: AB_218182 |
| Rabbit polyclonal anti-Ubiquitin | Cell Signaling Technology | E4I2J, 43124; RRID: AB_2799235 |
| Rat polyclonal anti-CTIP2 | Abcam | ab18465; RRID: AB_2064130 |
| Mouse monoclonal anti-BrdU | Millipore | MAB3424; RRID: AB_94851 |
| Rabbit polyclonal anti-Virma | ProteinTech | 25712-1-AP; RRID: AB_2880204 |
| Rabbit anti-Ubiquitin D9D5 (K48 linkage-specific) | Cell Signaling Technology | 8081; RRID: AB_10859893 |
| Rabbit polyclonal anti-beta III Tubulin | Abcam | ab18207; RRID: AB_444319 |
| Rabbit polyclonal anti-Cre | SySy | 257 003; RRID: AB_2619968 |
| Mouse monoclonal anti-ZO1 1A12 | Merck | 33-9100; RRID: AB_2533147 |
| Rabbit anti-Ki67 | Gift from Prim Singh | N/A |
| Mouse monoclonal anti 6X His-Tag | Abcam | ab18184; RRID: AB_444306 |
| Mouse anti-His, HIS.H8 | Abcam | ab18184; RRID: AB_444306 |
| Mouse anti-myc, 9B11 | Cell Signaling Technology | 2276; RRID: AB_2148465 |
| Rabbit anti-myc 71D10 | Cell Signaling Technology | 2278; RRID: AB_490778 |
| Rabbit anti-HA | Cell Signaling Technology | 3724; RRID: AB_1549585 |
| Rabbit anti-m6A | SySy | 202 003; RRID: AB_2279214 |
| Goat anti-EGFP | Rockland | 600-101-215; RRID: AB_218182 |
| **Secondary Antibodies** |  |  |
| Donkey anti-goat AlexaFluor-488 | Jackson ImmunoResearch | 705-545-003; RRID: AB_2340428 |
| Donkey anti-rabbit AlexaFluor-488 | Jackson ImmunoResearch | 711-545-152; RRID: AB_2313584 |
| Donkey anti-rabbit AlexaFluor-Cy3 | Jackson ImmunoResearch | 711-165-152; RRID: AB_2307443 |
| Donkey anti-rat AlexaFluor-Cy3 | Jackson ImmunoResearch | 712-165-153; RRID: AB_2340667 |
| Peroxidase IgG Fraction Monoclonal Mouse Anti-Rabbit IgG, light chain specific | Jackson ImmunoResearch | 211-032-171; RRID: AB_2339149 |
| Peroxidase AffiniPure Goat Anti-Mouse IgG, light chain specific | Jackson ImmunoResearch | 115-035-174; RRID: AB_2338512 |
| Donkey anti-Goat IgG (H+L) Cross-Adsorbed Secondary Antibody, Alexa Fluor 488 | Invitrogen | A11055; RRID: AB_2534102 |
| Donkey anti-Mouse IgG (H+L) Highly Cross-Adsorbed Secondary Antibody, Alexa Fluor 488 | Invitrogen | A21202; RRID: AB_141607 |
| Donkey anti-Rabbit IgG (H+L) Highly Cross-Adsorbed Secondary Antibody, Alexa Fluor 488 | Invitrogen | A21206; RRID: AB_2535792 |
| Donkey anti-Rabbit IgG (H+L) Highly Cross-Adsorbed Secondary Antibody, Alexa Fluor 555 | Invitrogen | A31572; RRID: AB_162543 |
| Donkey anti-mouse IgG (H+L) Highly Cross-Adsorbed Secondary Antibody, Alexa Fluor 555 | Invitrogen | A31570; RRID: AB_2536180 |
| Donkey anti-Mouse IgG (H+L) Highly Cross-Adsorbed Secondary Antibody, Alexa Fluor 647 | Invitrogen | A31571; RRID: AB_162542 |
| Donkey anti-Rat IgG (H+L) Highly Cross-Adsorbed Secondary Antibody, Alexa Fluor Plus 647 | Invitrogen | A48272; RRID: AB_2893138 |
| Donkey anti-Rabbit IgG (H+L) Highly Cross-Adsorbed Secondary Antibody, Alexa Fluor Plus 647 | Invitrogen | A31573; RRID: AB_2536183 |
| **Chemicals, peptides, and recombinant proteins** |  |  |
| NEM | Sigma-Aldrich | E3876 |
| BrdU | Abcam | ab142567 |
| STM2457 | Sigma-Aldrich | SML3360 |
| PMSF | Roche | 10837091001 |
| Benzonase | Merck | 70664-3 |
| 50x Complete Inhibitors | Sigma-Aldrich | 5892988001 |
| cOmplete protease inhibitor cocktail | Roche | 11873580001 |
| Ube3C HECT wild-type fragment | This study | N/A |
| Ube3C HECT C1051S fragment | This study | N/A |
| Ube3C HECT S845F fragment | This study | N/A |
| Ube3C HECT F996C fragment | This study | N/A |
| Coomassie Brilliant Blue | Neo-Biotech | NB-45-00078 |
| Invitrogen SilverXpress | Fisher Scientific | 10052662 |
| Protein G Sepharose beads | GE Healthcare | 17-0618-01 |
| Ni Sepharose 6 Fast Flow | Cytiva | 17531801 |
| GFP-Trap Agarose | ChromoTek | GTA |
| **Critical commercial assays** |  |  |
| BCA Protein Assay Kit | Thermo Scientific | 23225 |
| m6A Dot Blot Assay Kit | RayBiotech | DB-m6A |
| RNAscope Multiplex Fluorescent V2 Assay | ACDBio | 323100 |
| EpiQuik m6A RNA Methylation Quantification Kit (Colorimetric) | EpigenTek | P9005 |
| Control TUBE | Tebu-bio | UM0500M |
| K48-TUBE | Tebu-bio | UM407M |
| **Experimental models: Cell lines** |  |  |
| HEK293T | Leibniz Institute | RRID: CVCL_0063 |
| JM8A3.N1 | N/A | RRID: CVCL_J960 |
| TMOi001-A | Thermo Fisher Scientific | A18945, RRID: CVCL_RM92 |
| **Experimental models: Organisms/strains** |  |  |
| NMRI wild-type mouse strain | Charles Rivers and Janvier Labs | RRID: IMSR_TAC:nmri |
| *Ube3C*<sup>fl/fl</sup> mouse line | This study | N/A |
| *Emx1*<sup>Cre/+</sup>; *Ube3C*<sup>fl/fl</sup> mouse line | This study | N/A |
| *Emx1*-Cre driver | 40 |  |
| **Recombinant DNA** |  |  |
| pCAG-EGFP | Addgene | #14883 |
| pCX::myr-Venus | Gift from Anna-Katerina Hadjantonakis | N/A |
| pCAG-iCre | Addgene | #89573 |
| pCAG-EGFP-Ube3C | This study | N/A |
| pCAG-EGFP-Ube3C-C1051S | This paper | N/A |
| pCAG-EGFP-Ube3C-F996C | This paper | N/A |
| pCAG-EGFP-Ube3C-S845F | This paper | N/A |
| pRAI-myc-Ube3C | This paper | N/A |
| pRAI-myc-Ube3C-C1051S | This paper | N/A |
| pCAG-myc-Ube3C | This paper | N/A |
| pCAG-myc-Ube3C-C1051S | This paper | N/A |
| pCAG-myc-Ube3C-F996C | This paper | N/A |
| pCAG-myc-Ube3C-S845F | This paper | N/A |
| pCAG-HA-Cbll1 | This paper | N/A |
| pCAG | Addgene | #11160 |
| pCAG-EGFPiCre | Gift from Richard Huganir | N/A |
| pCAG-hypPBase | Gift from Jean Livet | N/A |
| pCAG-PB-Ube3C | This paper | N/A |
| pCAG-PB-EGFP | This paper | N/A |
| **Software and algorithms** |  |  |
| ImageJ | Imagej.net | RRID: SCR_003070 |
| GraphPad Prism | graphpad.com | RRID: SCR_002798 |
| Adobe Photoshop | adobe.com/products/photoshop.html | RRID: SCR_014199 |
| GSEA Open Source | https://bioinformatics.sdstate.edu/go/ | N/A |
| AlphaFold 3 | 74 | N/A |

N/A - not available

---

**Fig. S1.**
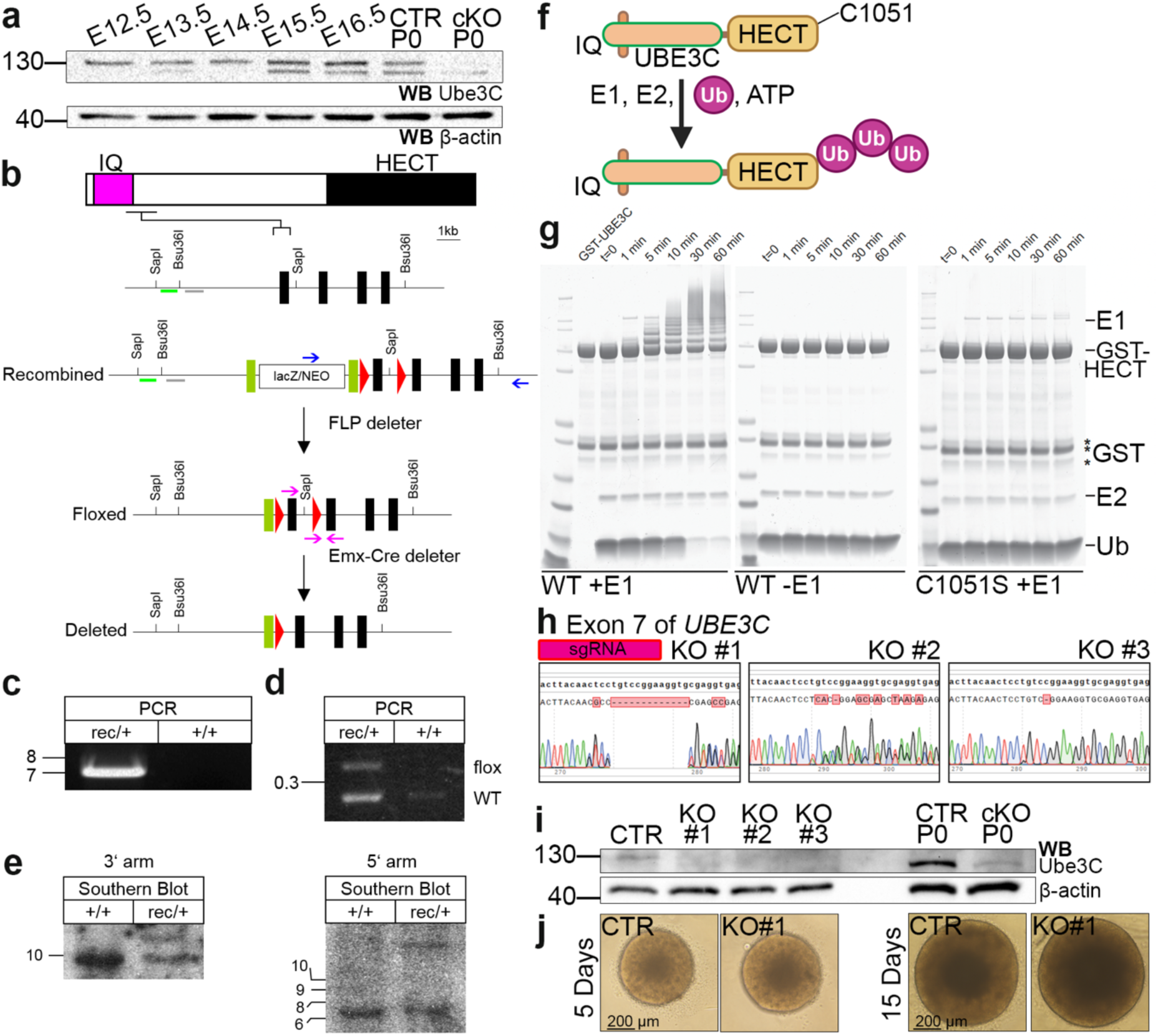
Related to the entire manuscript. Generation of genetic and biochemical tools to study Ube3C in the developing brain. (a) Western blotting using anti-Ube3C GeneTex N1N3 antibody in cortical lysates at indicated developmental age and from *Ube3C* ^f/f^ (CTR), and *Ube3C* ^f/f^; *Emx1*^Cre/+^ (cKO). E, embryonic day; P, postnatal day. CTR and cKO depict P0 cortical lysates from embryos of conditional mouse line. Raw Western blotting data used for representative figures are attached to this manuscript and available on Figshare data repository. (b) Gene targeting strategy of murine *Ube3C*. The domain structure of Ube3C, wild type, and targeted *Ube3C* alleles are shown. Exons, *loxP* sites, and FLP-recombinase target sites are symbolized as black, red, and green rectangles, respectively. Critical exon of *Ube3C* is flanked by two *loxP* sites. Mice positive for the recombined allele were crossed with FLP deleter and further crossed to *Emx1*-Cre-driver, leading to removal of exon 5 conditionally in the forebrain neurons and glia. (c, d, e) Validation of *Ube3C* gene targeting. +/+ wild type, rec/+ indicates heterozygous recombined mutant. DNA length indicators are in kilobase (kb). (c) The result of long-range PCR using primers depicted as blue arrows in (b). (d) Multiplex PCR using primers depicted as pink arrows in (b). PCR using genomic DNA from targeted ES cells yields two bands (left lane), and PCR using wild type ES cells genomic DNA as a template, results in one band. (e) Southern blotting analysis of genomic DNA purified from targeted ES cells. Genomic DNA isolated from control and ES cells carrying ’Recombined’ allele of *Ube3C* was digested with Bsu36I (3’) or SapI (5’) enzyme. The probes are indicated as coloured lines in (b). (f, g) A schematic of autoubiquitination activity of Ube3C. Representative Coomassie stained polyacrylamide gel showing in vitro autoubiquitylation assay depicted in (f) of recombinant GST-tagged HECT domain of Ube3C (GST-HECT). The reaction was composed of 0.05 μM UBA1 E1 (not added for a negative control), 1 μM UbcH5c E2, 3 μM GST-HECT-Ube3C or C1051S dominant negative variant, 15 μM Ub, and 4 mM ATP. (h) Results of Sanger sequencing using hg38 primers for electroporated iPSCs reveal successful targeting of exon 7 of *UBE3C*, generating deletions, leading to *UBE3C* KO from cells, validated by Western blotting, using abcam (ab243830) anti-Ube3C antibody (i). Reference sequence of exon 7 in bold. (j) Light microscopy of 5- and 15-days old organoids derived from isogenic CTR and *UBE3C* KO line.

**Fig. S2.**
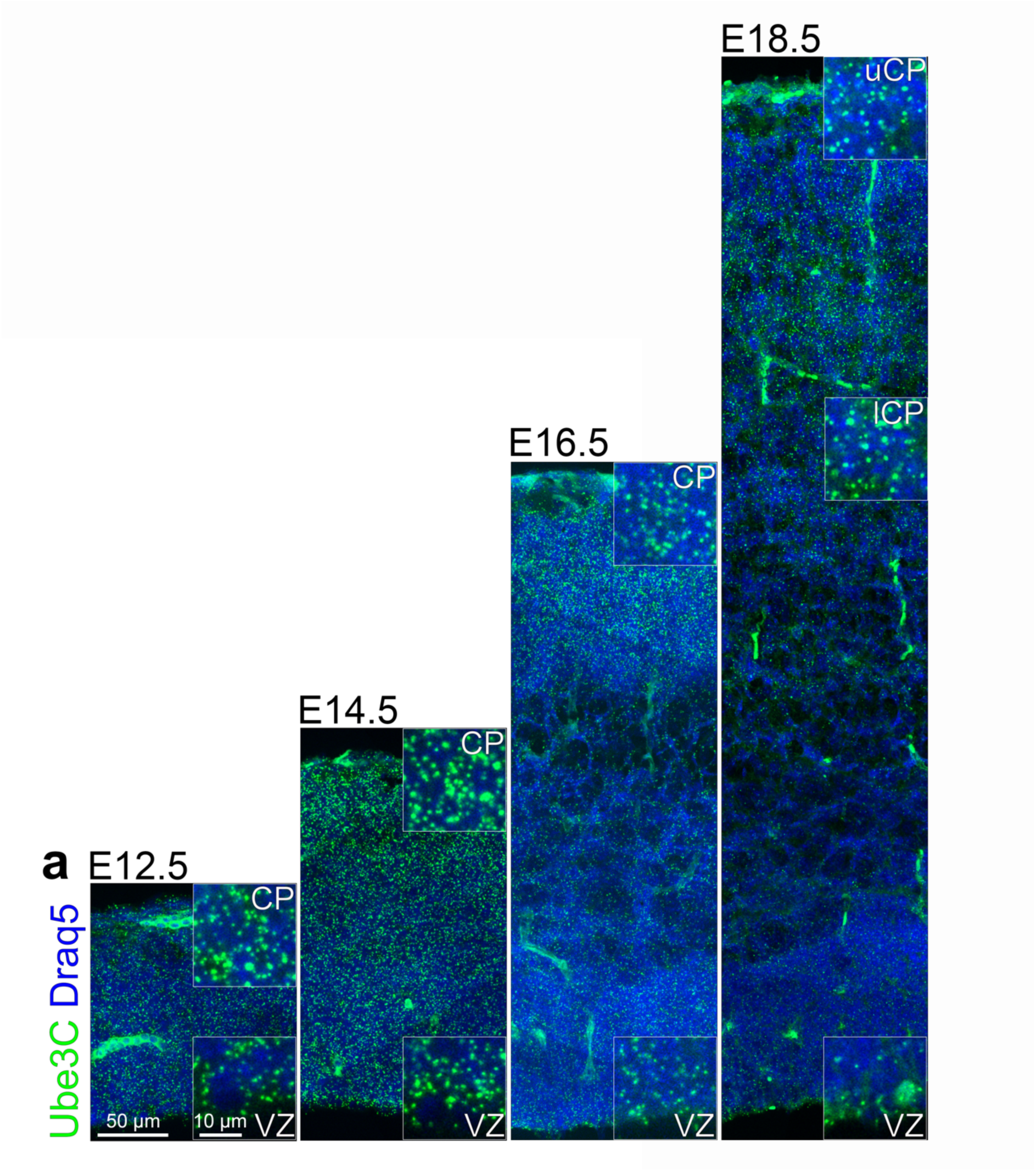
Related to the entire manuscript. Ube3C mRNA is expressed throughout the developing embryonic neocortex. (a) Coronal neocortical sections at indicated embryonic stages analyzed for Ube3C mRNA by FISH. Nuclei are stained with Draq5. Inset represent magnified indicated regions of the section. VZ, ventricular zone; CP, cortical plate; lCP, lower CP; uCP, upper CP.

**Fig. S3.**
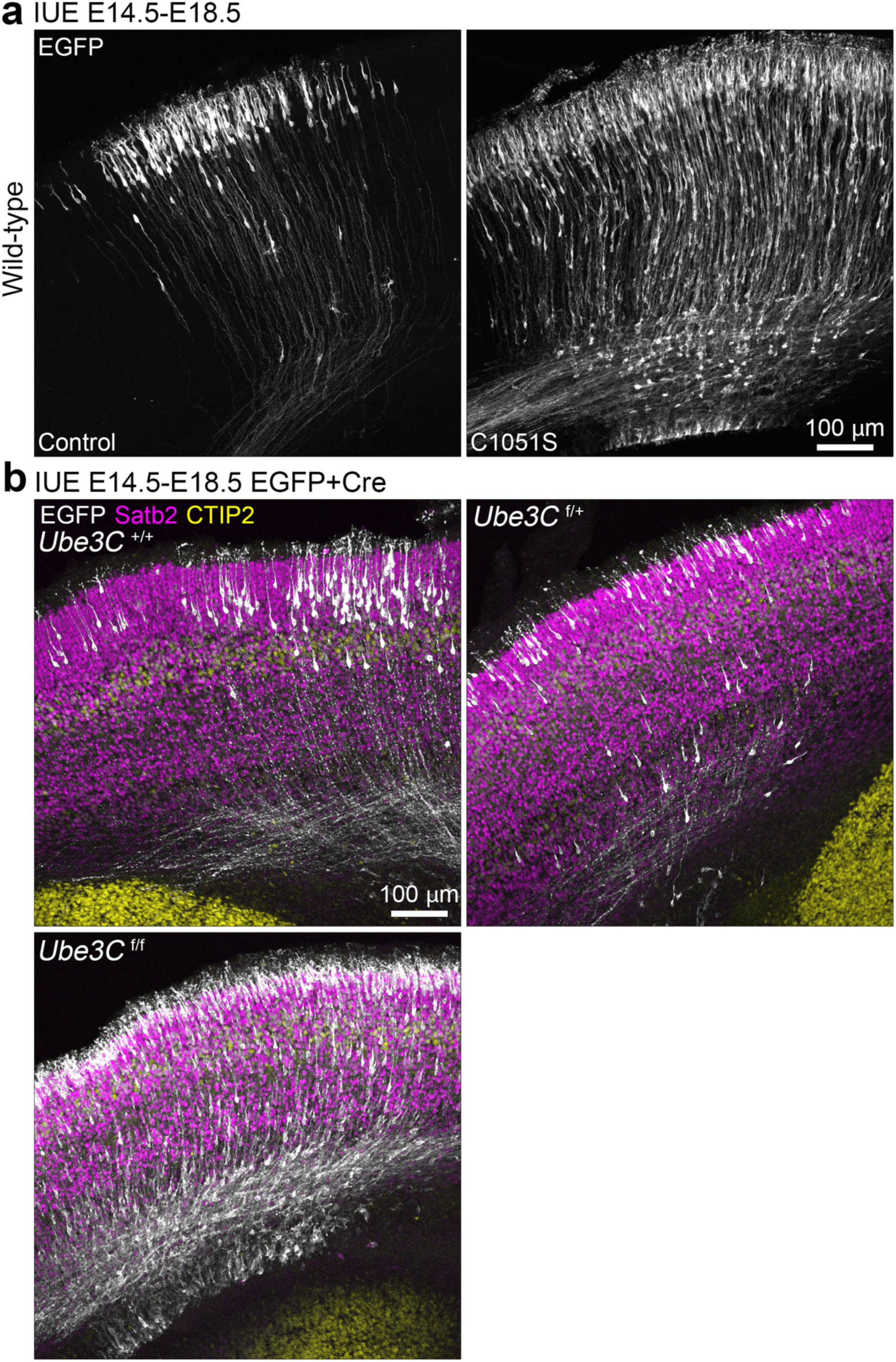
Related to Fig.1. Zoomed out images used for laminar positioning analysis for the anatomical context. (a) Representative images used for Fig. 1b. (b) Representative images used for Fig. 1e.

